# Kdm6b confers Tfdp1 with the competence to activate p53 signalling in regulating palatogenesis

**DOI:** 10.1101/2021.10.13.464272

**Authors:** Tingwei Guo, Xia Han, Jinzhi He, Jifan Feng, Junjun Jing, Eva Janečková, Jie Lei, Thach-Vu Ho, Jian Xu, Yang Chai

**Author notes:** Corresponding author: Yang Chai University Professor George and MaryLou Boone Chair in Craniofacial Biology Center for Craniofacial Molecular Biology University of Southern California 2250 Alcazar Street – CSA 103 Los Angeles, CA 90033.

## Abstract

Epigenetic regulation plays extensive roles in diseases and development. Disruption of epigenetic regulation not only increases the risk of cancer, but can also cause various developmental defects. However, it is still unclear how epigenetic regulators coordinate with tissue-specific regulatory factors during morphogenesis of specific organs. Using palatogenesis as a model, we reveal the functional significance of *Kdm6b*, a H3K27me3 demethylase, in regulating embryonic development. Our study shows that *Kdm6b* plays an essential role in neural crest development, and loss of *Kdm6b* disturbs p53 pathway-mediated activity, leading to complete cleft palate along with cell proliferation and differentiation defects. Furthermore, activity of H3K27me3 on the promoter of *p53* is precisely controlled by *Kdm6b*, and *Ezh2* in regulating p53 expression in cranial neural crest cells. More importantly, *Kdm6b* renders chromatin accessible to the transcription factor Tfdp1, which binds to the promoter of *p53* along with Kdm6b to specifically activate *p53* expression during palatogenesis. Collectively our results highlight the important role of the epigenetic regulator Kdm6b and how it cooperates with Tfdp1 to achieve its functional specificity in regulating *p53* expression, and further provide mechanistic insights into the epigenetic regulatory network during organogenesis.

## Introduction

Embryonic development is a highly complex self-assembly process during which precursor cells are coordinated to generate appropriate cell types and assemble them into well-defined structures, tissues, and organs (Shahbazi et al. 2016). During this process, precursor cells undergo extensive and rapid cell proliferation until they reach the point of exit from the cell cycle to differentiate into various cell lineages (Ruijtenberg and van den Heuvel 2016; Miermont et al. 2019). How these precursor cells modulate expression of different genes and proceed through diverse proliferation and differentiation processes is a very complex and interesting question. Growing evidence shows that epigenetic regulation, which includes mechanisms such as DNA methylation, histone modifications, chromatin accessibility, and higher-order organization of chromatin, provides the ability to modify gene expression and associated protein production in a cell type-specific manner, thus playing an essential role in achieving signaling specificity and in regulating cell fate during embryonic development (Hanna et al. 2018).

Among these various layers of epigenetic regulation, DNA methylation and histone methylation are the best-characterized and known to be key regulators of diverse cellular events (Bannister and Kouzarides 2011; Smith and Meissner 2013; Molina-Serrano et al. 2019). For example, methylation of lysine 27 on histone H3 (H3K27me) by methyltransferases is a feature of heterochromatin that renders it inaccessible to transcription factors, thus maintaining transcriptional repression, across many species (Wiles and Selker 2017). On the other hand, methylation of H3K4me3 found near the promoter region can couple with the NURF complex to increase chromatin accessibility for gene activation (Wysocka et al. 2006; Soares et al. 2017). Demethylation, which results from removing a methyl group, also plays important roles during development. For instance, demethylation of H3K4 is required for maintaining pluripotency in embryonic stem cells, and demethylases KDM6A and KDM6B are required for proper gene expression in mature T cells (Lessard and Crabtree 2010; Jambhekar et al. 2019). These studies clearly show that failure to maintain epigenomic integrity can cause deleterious consequences for embryonic development and adult tissue homeostasis (Henckel et al. 2007; Kim et al. 2009; Kang et al. 2019).

Palatogenesis is a complex process known to be regulated by multiple genetic regulatory mechanisms, including several signaling pathways (BMP, SHH, WNT, FGF, and TGFβ) and different transcription factors (such as *Msx1*, *Sox9*, *Lhx6/8*, *Dlx5*, *Shox2* and more) (Satokata and Maas 1994; Yu et al. 2005; Chai and Maxson 2006; Levi et al. 2006; Cobourne et al. 2009; Lee and Saint-Jeannet 2011; Nakamura et al. 2011; Bush and Jiang 2012; He and Chen 2012; Parada and Chai 2012; Xu et al. 2016; Reynolds et al. 2019). However, environmental effects can also contribute to orofacial defects, which lends further support to the notion that genetic factors are not sufficient to fully explain the etiology of many birth defects (Dixon et al. 2011; Roessler et al. 2012; Seelan et al. 2012). Furthermore, case studies have revealed that heterozygous mutation of a chromatin-remodeling factor, *SATB2*, and variation in DNA methylation can cause cleft palate in patients (Leoyklang et al. 2007; Chandrasekharan and Ramanathan 2014; Young et al. 2021). These cases have drawn our attention to the function of epigenetic regulation in palatogenesis.

The contribution of cranial neural crest cells (CNCCs) is critical to palate mesenchyme formation. Recently, studies have begun to address the role of epigenetic regulation in neural crest cell fates determination during development. For instance, homozygous loss of *Arid1a*, a subunit of SWI/SNF chromatin remodeling complex, in neural crest cells results in lethality in mice, associated with severe defects in the heart and craniofacial bones (Chandler and Magnuson 2016). In addition, both lysine methyltransferase *Kmt2a* and demethylase *Kdm6a* are essential for cardiac and neural crest development (Shpargel et al. 2017; Sen et al. 2020). However, how these epigenetic changes lead to tissue-specific response during neural crest fate determination remain to be elucidated.

In this study, using palatogenesis as a model we investigated the functional significance of the demethylase *Kdm6b* in regulating the fate of CNCCs during palatogenesis. We have discovered that loss of *Kdm6b* in CNC-derived cells results in complete cleft palate along with soft palate muscle defects. We also found cell proliferation and differentiation defects of CNC-derived cells in *Kdm6b* mutant mice. More importantly, our study shows that the level of H3K27me3 on the promoter of *p53* is precisely controlled by *Kdm6b*, and *Ezh2* in regulating expression of *p53* (also known as *Trp53*). Furthermore, the transcription factor Tfdp1 binds to the promoter of *p53* along with Kdm6b to specifically activate the expression of this tumor suppressor gene. Our study highlights the importance of epigenetic regulation on cell fate decision and its function in regulating activity of *p53* in CNC-derived cells during organogenesis.

## Results

### Loss of *Kdm6b* in CNC-derived cells results in craniofacial malformations

Previous research has shown that the X-chromosome-linked H3K27 demethylase Kdm6a is indispensable for neural crest cell differentiation and viability as it establishes appropriate chromatin structure (Schwarz et al. 2014; Shpargel et al. 2017). However, we do not yet have a comprehensive understanding of the roles of two other members of the Kdm6 family, *Kdm6b* and *Uty*, in regulating CNCCs during craniofacial development. More importantly, we have yet to understand how demethylase achieves its functional specificity in regulating downstream target genes. In order to elucidate the functions of *Kdm6b* and *Uty*, we first evaluated the expression patterns of Kdm6 family members in the palatal region (Figure 1-figure supplement 1A-F). We found that, of these, *Kdm6b* is more abundantly expressed than *Kdm6a* and *Uty* in both palate mesenchymal and epithelial cells, which indicated it might play a critical role in regulating palatogenesis.

To investigate the tissue-specific function of *Kdm6b* during craniofacial development, we generated *Wnt1-Cre;Kdm6b^fl/fl^* and *K14-Cre;Kdm6b^fl/fl^* mice to specifically target the deletion of *Kdm6b* in CNC-derived and epithelial cells, respectively. Loss of *Kdm6b* in CNC-derived cells resulted in complete cleft palate in *Wnt1-Cre;Kdm6b^fl/fl^* mice (90% phenotype penetrance, N = 7) and postnatal lethality at newborn stage (100% phenotype penetrance, N = 7) without interrupting expression of other Kdm6 family members (Figure 1-figure supplement 1A-N and Figure 1A-B). To evaluate when *Kdm6b* was inactivated in the CNC-derived cells, we also investigated the expression of *Kdm6b* at E9.5, well prior to the formation of the palate primordium, and found that *Kdm6b* was efficiently inactivated in the CNC-derived cells at this stage (Figure 1-figure supplement 1O-P). Interestingly, loss of *Kdm6b* in epithelial cells did not lead to obvious defects in the craniofacial region in *K14-Cre;Kdm6b^fl/fl^* mice (Figure 1-figure supplement 2A-H). These results emphasized that *Kdm6b* is specifically required in CNC-derived cells during palatogenesis. CT images also confirmed the complete cleft palate phenotype and revealed that the most severe defects in the palatal region of *Wnt1-Cre;Kdm6b^fl/fl^* mice were hypoplastic palatine processes of the maxilla and palatine bones (Figure 1C-D). Except for a minor flattened skull, other CNC-derived bones did not show significant differences between control and *Wnt1-Cre;Kdm6b^fl/fl^* mice (Figure 1-figure supplement 2I-N). To evaluate the phenotype in more detail, we performed histological analysis and found that although the palatal shelves were able to elevate, the maxilla and palatine bones, as well as the palate stromal mesenchyme and soft palate muscles, failed to grow towards the midline in *Wnt1-Cre;Kdm6b^fl/fl^* mice (Figure 1G-R and Figure 1-figure supplement 3A-P). Furthermore, in the posterior soft palate region, *Wnt1-Cre;Kdm6b^fl/fl^* mice also showed morphological defects related to the orientation of muscle fibers and the pterygoid plate (Figure 1-figure supplement 3A-X). However, since the soft palate forms subsequent to the hard palate, it is difficult to identify whether the soft palatal muscle phenotype is a primary defect or a consequence resulting from an anterior cleft. Therefore, we focused on the anterior hard palate for further investigation. Collectively, these data indicate that mesenchymal *Kdm6b* is indispensable for craniofacial development and plays an essential role during palatogenesis.

**Figure 1.**
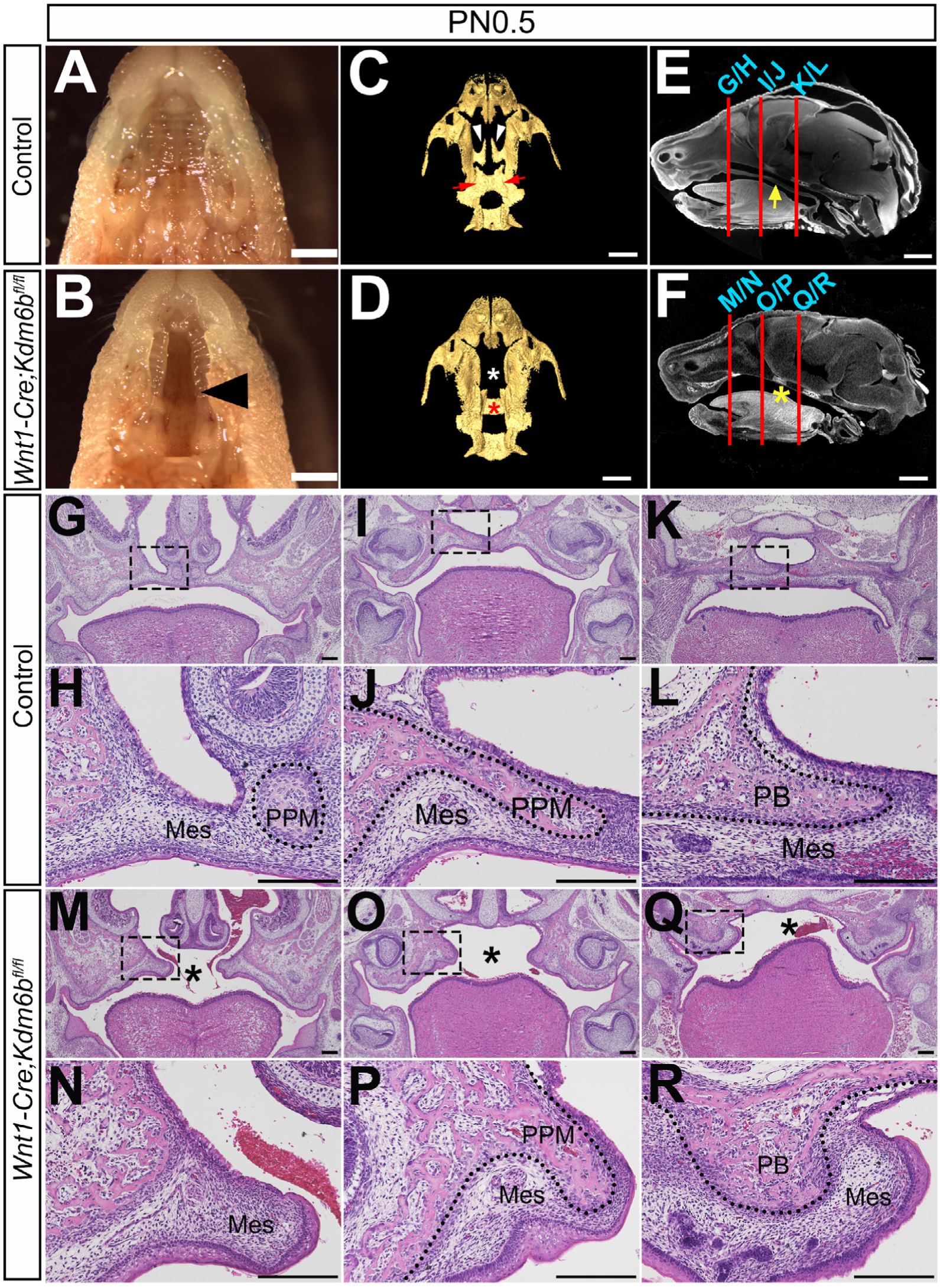
Loss of *Kdm6b* results in cleft palate. (A-B) Whole-mount oral view shows complete cleft palate phenotype in *Wnt1-Cre;Kdm6b^fl/fl^* mice. Arrowhead in B indicates the cleft palate. Scale bar: 2 mm. (C-D) CT imaging reveals that the palatine process of the maxilla and palatine bone are missing in *Wnt1-Cre;Kdm6b^fl/fl^* mice. White arrowheads in C indicates palatine process of maxilla in control mice, and red arrows indicates palatine bone. White asterisk in D indicates missing palatine process of maxilla in *Wnt1-Cre;Kdm6b^fl/fl^* mice and red asterisk in D indicates the missing palatine bone in *Kdm6b* mutant mice. Scale bars: 1 mm. (E-F) Sagittal views of CT images demonstrate the locations of HE sections in G-R. Red lines indicate the locations of sections. Yellow arrow in E indicates palatal shelf and yellow asterisk in F indicates cleft. Scale bars: 1 mm. (G-R) Histological analysis of control and *Wnt1-Cre;Kdm6b^fl/fl^* mice. H, J, L, N, P, and R are magnified images of boxes in G, I, K, M, O, and Q, respectively. Asterisks in M, O, and Q indicate cleft in *Kdm6b* mutant mice. Scale bar: 200 µm. Mes: mesenchyme; PPM: palatine process of maxilla; PB: palatine bone.

### *Kdm6b* is critical for proliferation and differentiation of CNC-derived palatal mesenchymal cells

During craniofacial development, CNCCs migrate ventro-laterally and populate the branchial arches to give rise to distinct mesenchymal structures in the head and neck, such as the palate. Failure of CNCCs to populate pharyngeal arches causes craniofacial defects (Noden 1983; Noden 1991; Trainor and Krumlauf 2000; Cordero et al. 2011). To determine whether *Kdm6b* mutant CNCCs successfully populate the first pharyngeal arch, which gives rise to the palatal shelves, we generated tdTomato reporter mice and collected samples at E10.5. The results showed that CNCCs migration was not adversely affected in *Kdm6b* mutant mice (Figure 2-figure supplement 4A-B). Then we evaluated the process of palatogenesis at different embryonic stages, and found that the cleft palate phenotype emerged as early as E14.5 in *Wnt1-Cre;Kdm6b^fl/fl^* mice (Figure 2-figure supplement 4C-D). These data established that *Kdm6b* is not essential for CNCCs entering the pharyngeal arch but is specifically required in regulating post-migratory CNC-derived cells.

Because cell proliferation defects in CNC-derived cells frequently lead to craniofacial defects, we tested whether loss of *Kdm6b* can affect cell proliferation using EdU labeling. After 2 hours of EdU labeling, we found that the number of cells positively stained with EdU were significantly increased in the CNC-derived palatal mesenchyme in *Wnt1-Cre;Kdm6b^fl/fl^* mice compared to controls (Figure 2A-C). In addition, after 48 hours of EdU labeling, we found that the number of Ki67 and EdU double-positive cells was significantly increased in *Wnt1-Cre;Kdm6b^fl/fl^* mice (Figure 2D-H). These results indicated that loss of *Kdm6b* in CNC-derived cells resulted in more cells remaining in the cell cycle and actively proliferating, which further led to hyperproliferation of mesenchymal cells in the palatal region of *Wnt1-Cre;Kdm6b^fl/fl^* mice. Meanwhile, increased cell death in the palatal mesenchyme was observed in *Wnt1-Cre;Kdm6b^fl/fl^* mice compared to controls, based on TUNEL staining analysis (Figure 2-figure supplement 4E-H).

**Figure 2.**
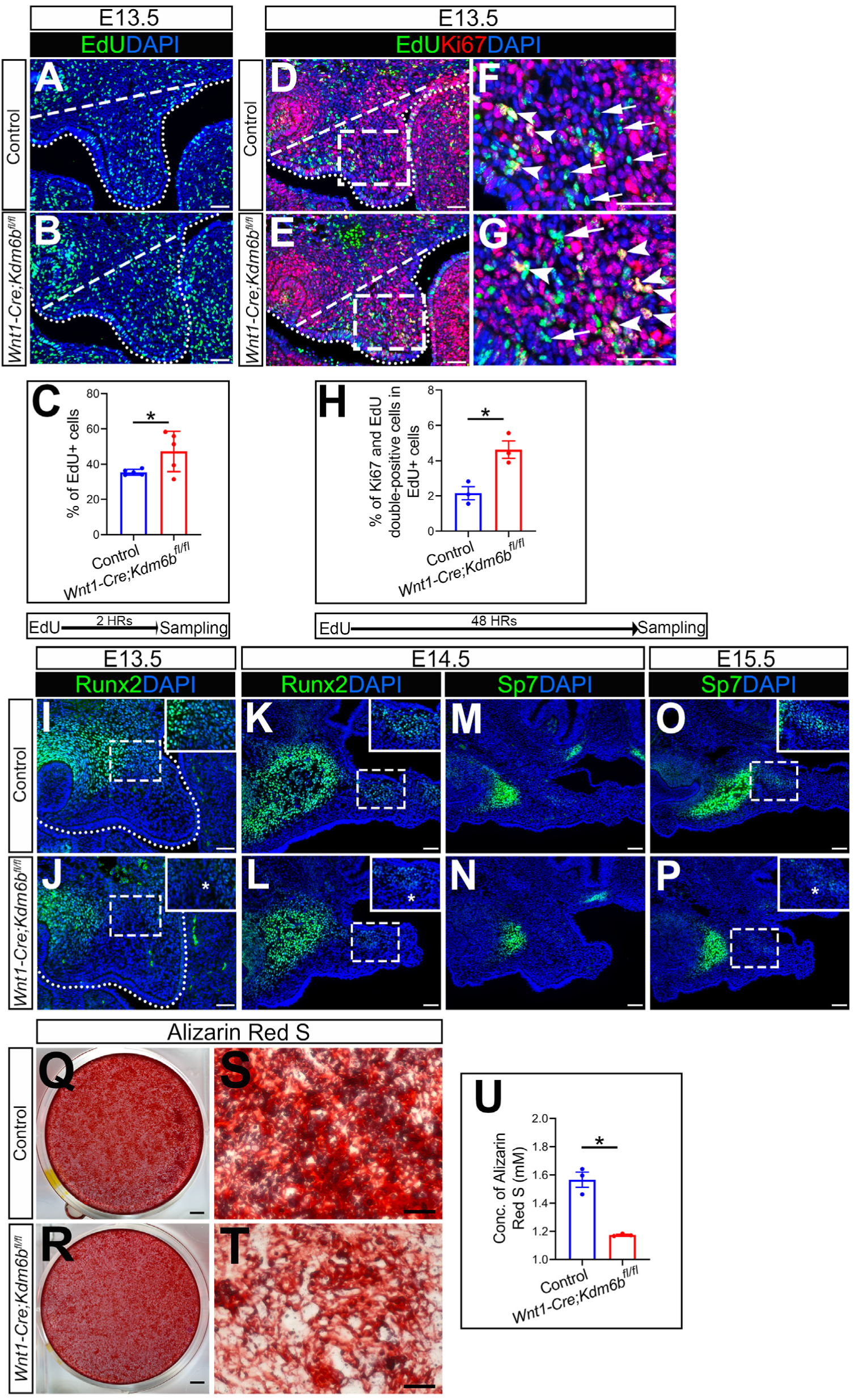
*Kdm6b* is critical for proliferation and differentiation of CNC-derived palatal mesenchyme cells. (A-B) Immunostaing of EdU at E13.5 after 2 hours of EdU labeling. Dotted lines indicate palatal shelf region. Dashed lines indicate the palatal region used for quantification in C. Scale bar: 50 µm. (C) Quantification of EdU+ cells represented in A and B. (D-G) Co-localization of EdU and Ki67 at E13.5 after 48 hours of EdU labeling. Dotted lines indicate palatal shelf region. Dashed lines indicate the palatal region used for quantification in H. F and G are magnified images of boxes in D and E. Arrows in F and G indicate representative cells that are only EdU+, while arrowheads indicate representative cells that are positive for both EdU and Ki67. Scale bar: 50 µm. (H) Quantification of EdU and Ki67 double-positive cells represented in D and E. (I-L) Immunostaining of Runx2 at indicated stages. Insets are higher magnification images of boxes in I-L. Asterisks in J and L indicate decreased Runx2+ cells observed in *Wnt1-Cre;Kdm6b^fl/fl^* mice. Scale bar: 50 µm. (M-P) Immunostaining of Sp7 at indicated stages. Insets are higher magnification images of boxes in O and P. Asterisk in P indicates decreased Sp7+ cells observed in *Wnt1-Cre;Kdm6b^fl/fl^* mice. Scale bar: 50 µm. (Q-U) Osteogenenic differentiation assay using Alizarin red S staining. U is the quantification result of Alizarin red S staining represented in Q and R. Scale bars: 2 mm in Q and R; 200 µm in S amd T. Figure 2-Source data 1 for figure 2C Figure 2-Source data 2 for figure 2H Figure 2-source data 3 for figure 2U

Typically, cell proliferation and differentiation are inversely correlated. Differentiation of precursor cells is generally associated with arrested proliferation and permanently exiting the cell cycle (Ruijtenberg and van den Heuvel 2016). To test whether cell differentiation was affected in the CNC-derived palatal mesenchyme in *Wnt1-Cre;Kdm6b^fl/fl^* mice, we examined the distribution of early osteogenesis markers Runx2 and later osteogenesis marker Sp7 in the palatal region from E13.5 to E15.5 (Figure 2I-P). There was a decrease in the number of Runx2+ cells in the palatal mesenchyme at both E13.5 and E14.5 in *Wnt1-Cre;Kdm6b^fl/fl^* mice in comparison to the control (Figure 2I-L). In addition, Sp7+ cells were also decreased in *Wnt1-Cre;Kdm6b^fl/fl^* mice at both E14.5 and E15.5 (Figure 2M-P). Furthermore, when we induced osteogenic differentiation in palatal mesenchymal cells from E13.5 embryos for three weeks, we found that cells from *Wnt1-Cre;Kdm6b^fl/fl^* mice showed much less calcium deposition than cells from control mice, indicating a reduction in osteogenic potential in cells from *Kdm6b* mutant mice (Figure 2Q-U). These results indicated that *Kdm6b* was indispensable for maintaining normal proliferation and differentiation of CNC-derived cells.

### Loss of *Kdm6b* in CNC-derived cells disturbs p53 pathway-mediated activity

In order to identify the downstream targets of *Kdm6b* in the palatal mesenchyme, we performed RNA-seq analysis of palatal tissue at E12.5. The results showed that more genes were downregulated than upregulated in the palatal mesenchyme in *Wnt1-Cre;Kdm6b^fl/fl^* mice (Figure 3A), which is consistent with the function of *Kdm6b* in removing the repressive mark H3K27me3. We further used Ingenuity Pathway Analysis (IPA) and Gene Ontology (GO) analysis to analyze the pathways that were most disturbed in the palatal mesenchyme in *Kdm6b* mutant mice. Surprisingly, both analyses indicated that pathways involving *p53* might be disturbed in the palatal mesenchyme in *Wnt1-Cre;Kdm6b^fl/fl^* mice (Figure 3B-C).

**Figure 3.**
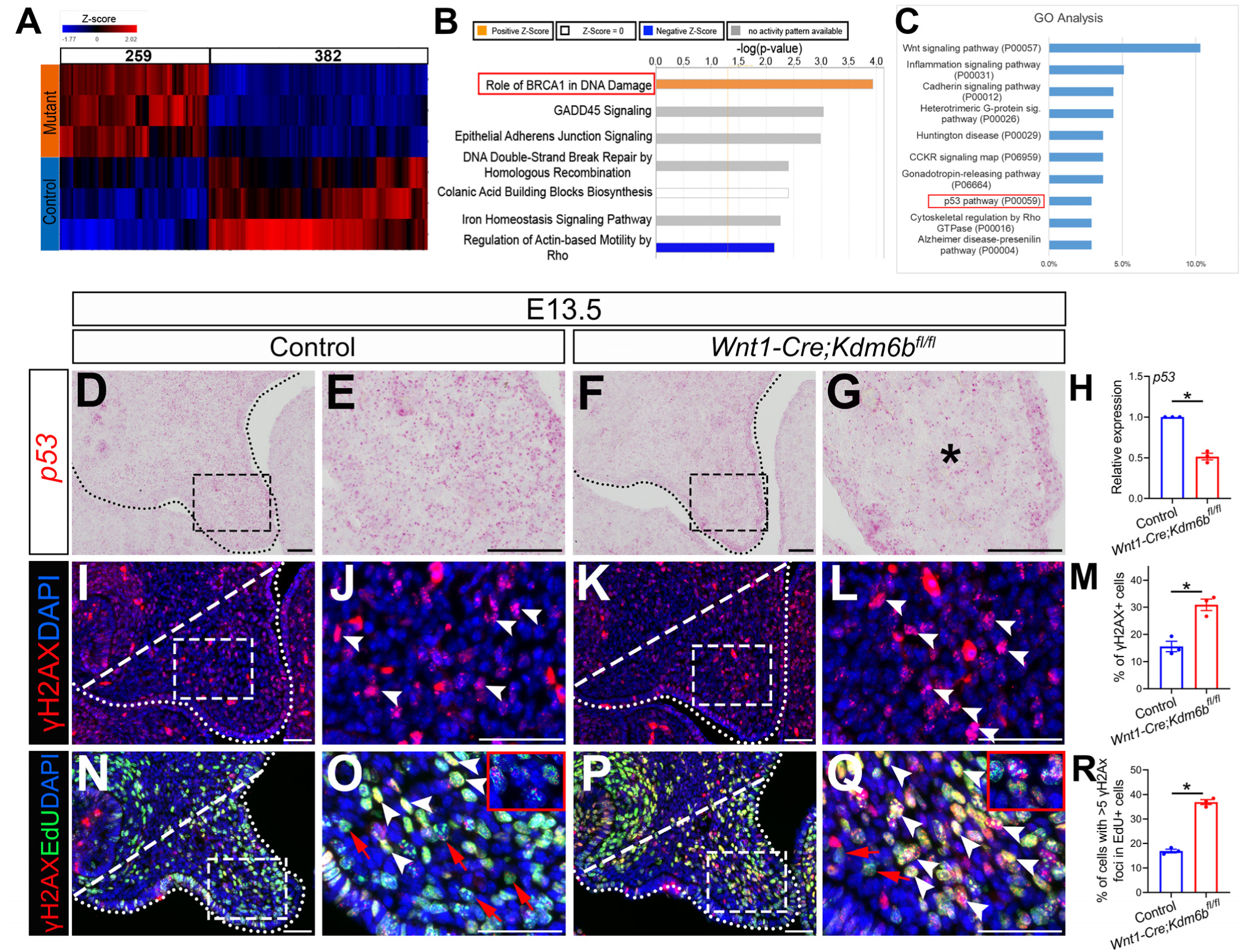
p53 signaling pathway is disturbed in *Wnt1-Cre;Kdm6b^fl/fl^* mice. (A) Bulk RNA-seq result of palatal tissues collected at E12.5 is represented in heatmap. Differentially expressed genes are selected using P < 0.05 and fold change < -1.2 or > 1.2. (B) IPA analysis using of bulk RNA-seq result. Red box indicates the top upregulated pathway observed in *Wnt1-Cre;Kdm6b^fl/fl^* sample. (C) GO analysis using bulk RNA-seq result. Red box indicates p53 signaling is one of the top 10 pathways. Y-axis shows the percentage of genes hit against total number of pathways hit. (D-G) Expression of *p53* at E13.5 using RNAscope *in situ* hybridization. Dotted lines in D and F indicate palatal shelf. E and G are magnified images of boxes in D and F. Asterisk in G indicates decreased expression of *p53* observed in *Wnt1-Cre;Kdm6b^fl/fl^* mice. Scale bar: 50 µm. (H) RT-qPCR quantification of *p53* in palatal tissues collected at E13.5. Asterisk indicates P < 0.05. (I-L) Immunostaining of γH2AX at E13.5. Dotted lines in I and K indicate palatal shelf and dashed lines indicate quantification area. J and L are magnified images of boxes in I and K, respectively. Arrowheads in J and L indicate representative γH2AX+ cells. Scale bar: 50 µm. (M) Quantification of γH2AX+ cells represented in I and K. Asterisk indicates P < 0.05. (N-Q) Co-localization of EdU and γH2AX at E13.5 after 2 hours of EdU labeling. Dotted lines in N and P indicate palatal shelf region, while dashed lines indicate the palatal region used for quantification in R. O and Q are magnified images of boxes in N and P. Red arrows in O and Q indicate representative EdU+ cells with less than 5 γH2AX foci, while white arrowheads indicate representative cells that are positive for EdU and with >5 γH2AX foci. Scale bar: 50 µm. (R) Quantification of EdU+ cells with >5 γH2AX foci represented in N and P. Asterisk indicates P < 0.05. Figure 3-Source data 1 for figure 3H Figure 3-Source data 2 for figure 3M Figure 3-Source data 3 for figure 3R

The tumor suppressor p53 plays prominent roles in regulating DNA damage response, including arresting cell growth for DNA repair, directing cellular senescence, and activating apoptosis (Mijit et al. 2020). Mutation of *p53* is a major cause of cancer development (Williams and Schumacher 2016). Previous research has shown that homozygous *p53* mutant mice exhibit craniofacial defects with complete cleft palate, while inappropriate activation of *p53* during embryogenesis also causes developmental defects including craniofacial abnormalities (Tateossian et al. 2015; Bowen et al. 2019). These results suggest that precise dosage of *p53* is indispensable for craniofacial development. We analyzed expression of *p53* in our samples and found that it significantly decreased in the palatal region of the *Kdm6b* mutant mice (Figure 3D-H). These results indicate that *Kdm6b* plays an important role in regulating the p53 pathway in the CNC-derived mesenchyme during palatogenesis. To further evaluate the consequence of downregulated *p53* in *Wnt1-Cre;Kdm6b^fl/fl^* mice, we assessed DNA damage, which are tightly related to the function of *p53*, in our study. We found that DNA damage increased, as indicated by γH2AX expression, in the palatal mesenchyme in *Wnt1-Cre;Kdm6b^fl/fl^* mice (Figure 3I-M). More importantly, we observed significantly increased γH2AX foci in the EdU+ cells of *Wnt1-Cre;Kdm6b^fl/fl^* palatal mesenchyme (Figure 3N-R). These data indicated that actively proliferating cells in *Wnt1-Cre;Kdm6b^fl/fl^* mice experienced more severe DNA damage compared to those in the control mice, which might be the result of replication stress caused by the hyperproliferation we observed in *Wnt1-Cre;Kdm6b^fl/fl^* mice. Collectively, these results suggest that *p53*’s function in DNA damage response is impaired in *Wnt1-Cre;Kdm6b^fl/fl^* mice. Furthermore, without appropriate activation of *p53*, cells of the palatal mesenchyme escaped from proper cell cycle arrest for DNA damage repair, which resulted in increased DNA damage in the *Kdm6b* mutant mice.

### Altered *p53* expression is responsible for the developmental defects in *Wnt1-Cre;Kdm6b^fl/fl^* mice

To further test whether downregulated expression of *p53* is a key factor in the developmental defects we observed in *Wnt1-Cre;Kdm6b^fl/fl^* mice, we transfected palatal mesenchymal cells from control mice with siRNA to knock down *p53*. qPCR revealed that expression of *p53* was significantly decreased in the cells treated with siRNA after three days (Figure 4-figure supplement 5A). At the same time, the group transfected with siRNA for *p53* showed a significant increase in EdU+ cells (Figure 4A-C). In addition, significantly increased γH2AX+ cells were also observed in the group transfected with siRNA for *p53* (Figure 4D-F). These data suggested that downregulated expression of *p53* in the palatal mesenchymal cells is a key factor that led to the hyperproliferation and increased DNA damage we observed in *Wnt1-Cre;Kdm6b^fl/fl^* mice. Furthermore, both expression of *Runx2* and *Sp7* were also significantly reduced in the palatal mesenchymal cells transfected with siRNA for *p53* (Figure 4-figure supplement 5B-C), which indicated that downregulated expression of *p53* in the palatal mesenchymal cells resulted in differentiation defects, which were also observed in *Wnt1-Cre;Kdm6b^fl/fl^* mice.

**Figure 4.**
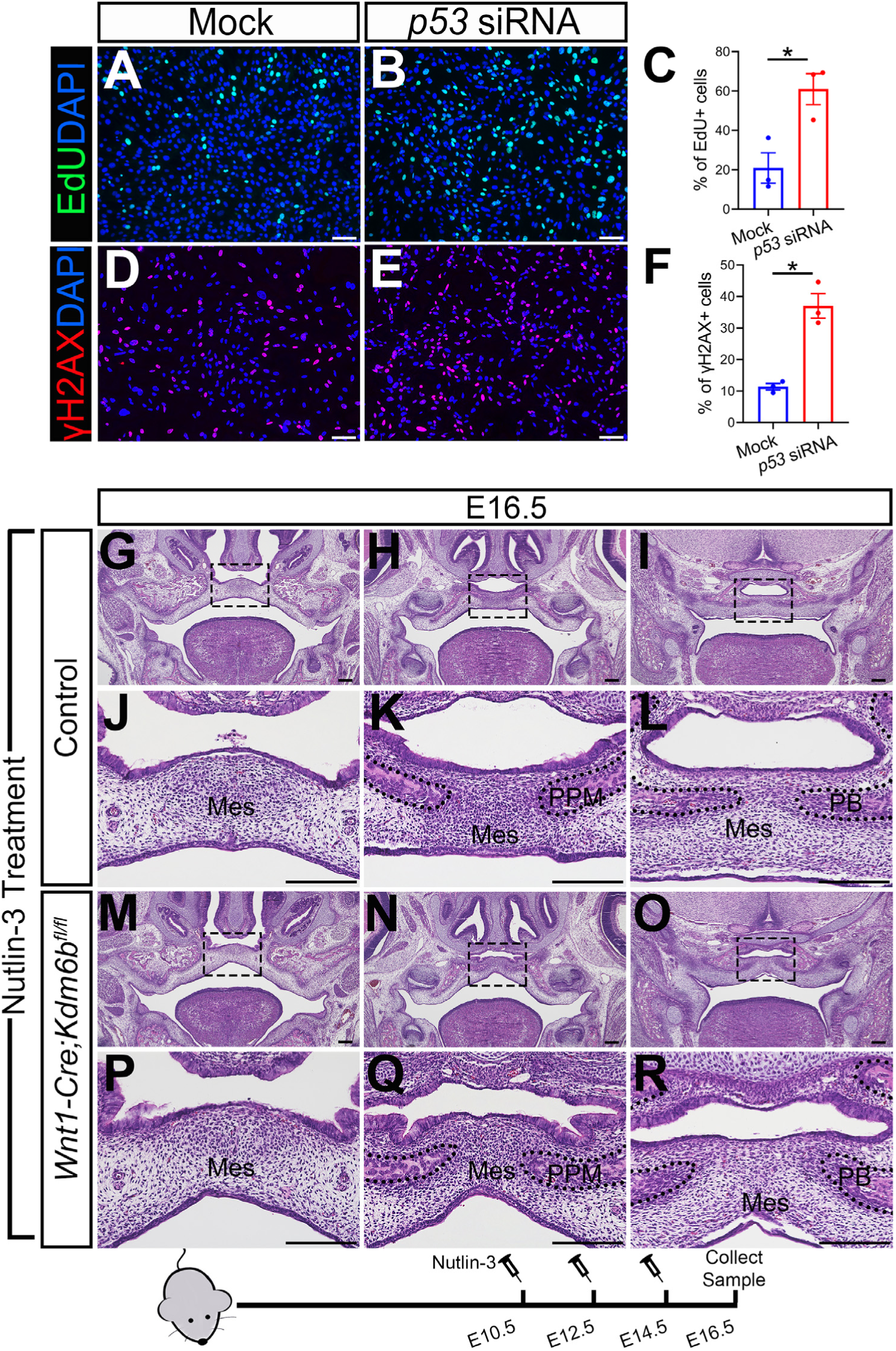
Altered *p53* expression is responsible for the developmental defects in *Wnt1-Cre;Kdm6b^fl/fl^* mice. (A-C) Cells collected from E13.5 palatal tissue are transfected with siRNA to knock down expression of *p53*. Cell proliferation is evaluated using EdU labeling 3 days after transfection. A and B show proliferation of cells assessed by EdU labeling. Difference in EdU+ cells between mock- and siRNA-transfected groups is quantified in C. Scale bar: 100 µm. Asterisk indicates P < 0.05. (D-F) Cells collected from E13.5 palatal tissue are transfected with siRNA to knock down expression of *p53*. DNA damage is evaluated using γH2AX 3 days after transfection. D and E show γH2AX+ cells. Difference in γH2AX+ cells between mock- and siRNA-transfected groups is quantified in F. Scale bar: 100 µm. Asterisk indicates P < 0.05. (G-R) Histological analysis of control and *Wnt1-Cre;Kdm6b^fl/fl^* mice treated with Nutlin-3. J, K, L, P, Q, and R are magnified images of boxes in G, H, I, M, N, and O, respectively. Scale bar: 200 µm. Mes: mesenchyme; PPM: palatine process of maxilla; PB: palatine bone. Figure 4-Source data 1 for figure 4C Figure 4-Source data 2 for figure 4F

To further investigate the function of *p53* in *Wnt1-Cre;Kdm6b^fl/fl^* mice, we tried to increase p53 in *Kdm6b* mutant mice using available small molecules. Previous research showed that MDM2, a ubiquitin ligase, specifically targets p53 for degradation and there is increased p53 activity in *Mdm2* mutant mice, which exhibit a range of developmental defects (Arya et al. 2010; Bowen and Attardi 2019; Bowen et al. 2019). Nutlin-3, an MDM2 inhibitor that can specifically interrupt interaction between MDM2 and p53, increases p53 in mouse primary neural stem progenitor cells and rescues neurogenic deficits in *Fmr1* KO mice (Li et al. 2016). We treated pregnant mice with Nutlin-3 at a dosage based on their body weight at E10.5, E12.5 and E14.5 of pregnancy and then collected samples at E16.5 for analysis. To assess the potential influence of the solvent used to dissolve Nutlin-3 (10% DMSO in corn oil), we also treated mice with 10% DMSO in corn oil at the same embryonic stages. None of the *Kdm6b* mutant mice were rescued after this treatment (N=3) (Figure 4-figure supplement 5D-E). In contrast, Nutlin-3 treatment successfully rescued the cleft palate observed in *Wnt1-Cre;Kdm6b^fl/fl^* mice (N = 5) (Figure 4G-R). Western blot showed that the protein level of p53 was successfully restored in the Nutlin-3-treated group (Figure 4-figure supplement 5F). This result further revealed that downregulation of *p53* in *Wnt1-Cre;Kdm6b^fl/fl^* mice plays an essential role in the palatal defects and genetic interaction between *Kdm6b* and *p53*, and that it is important for the development of post-migratory CNCCs.

### Level of H3K27me3 is precisely regulated by *Kdm6b* and *Ezh2* during palatogenesis

The lysine-specific demethylase Kdm6b is able to activate gene expression via removing the H3K27me3 repressive mark (Jiang et al. 2013). To investigate whether *Kdm6b* regulates the expression of *p53* through modifying the level of H3K27me3, we first examined the status of H3K27me3 in our samples and found that loss of *Kdm6b* in CNC-derived cells resulted in accumulation of H3K27me3 in the nucleus of CNC-derived palatal mesenchymal cells (Figure 5A-D). Furthermore, immunoblotting revealed that the level of H3K27me3 was increased in the palatal region of *Kdm6b* mutant mice (Figure 5E). Since the level of H3K27me3 can also be modified by the methyltranferases Ezh1 and Ezh2, we further evaluated whether expression of Ezh1 and Ezh2 was affected in the palatal region. We found no obvious differences in either the distribution of *Ezh1+* cells or the Ezh1 protein level between control and *Kdm6b* mutant mice (Figure 5F-J). Similarly, no dramatic changes were observed in either the distribution of Ezh2+ cells or the Ezh2 protein level between control and *Kdm6b* mutant mice (Figure 5K-O). These results indicated that increased H3K27me3 in *Wnt1-Cre;Kdm6b^fl/fl^* mice was mainly caused by loss of *Kdm6b* in CNC-derived cells. However, we did notice a broader contribution and stronger signal of Ezh2 than Ezh1 in the CNC-derived palatal mesenchyme. To investigate whether an increase of H3K27me3 in the CNC-derived cells caused the cleft phenotype we observed in *Wnt1-Cre;Kdm6b^fl/fl^* mice, we generated *Wnt1-Cre;Kdm6b^fl/fl^;Ezh2^fl/+^* mice and assessed the level of H3K27me3 in this model. In *Wnt1-Cre;Kdm6b^fl/fl^;Ezh2^fl/+^* mice, we observed a rescue of the abnormal accumulation of H3K27me3 (Figure 5P-V). More importantly, haploinsufficiency of *Ezh2* in this model successfully rescued the cleft palate phenotype observed in *Wnt1-Cre;Kdm6b^fl/fl^* mice (Figure 6A-O) with 70% efficiency (N = 10). CT scanning showed that both the palatine processes of the maxilla and palatine bone were restored in the *Wnt1-Cre;Kdm6b^fl/fl^;Ezh2^fl/+^* mice (Figure 6A-C). Both bone and palatal mesenchymal tissue were rescued in *Wnt1-Cre;Kdm6b^fl/fl^;Ezh2^fl/+^* mice (Figure 6D-O). These results suggested that an antagonistic interaction between the histone demethylase Kdm6b and methyltransferase Ezh2 that modulates H3K27me3 is essential for palatogenesis.

**Figure 5.**
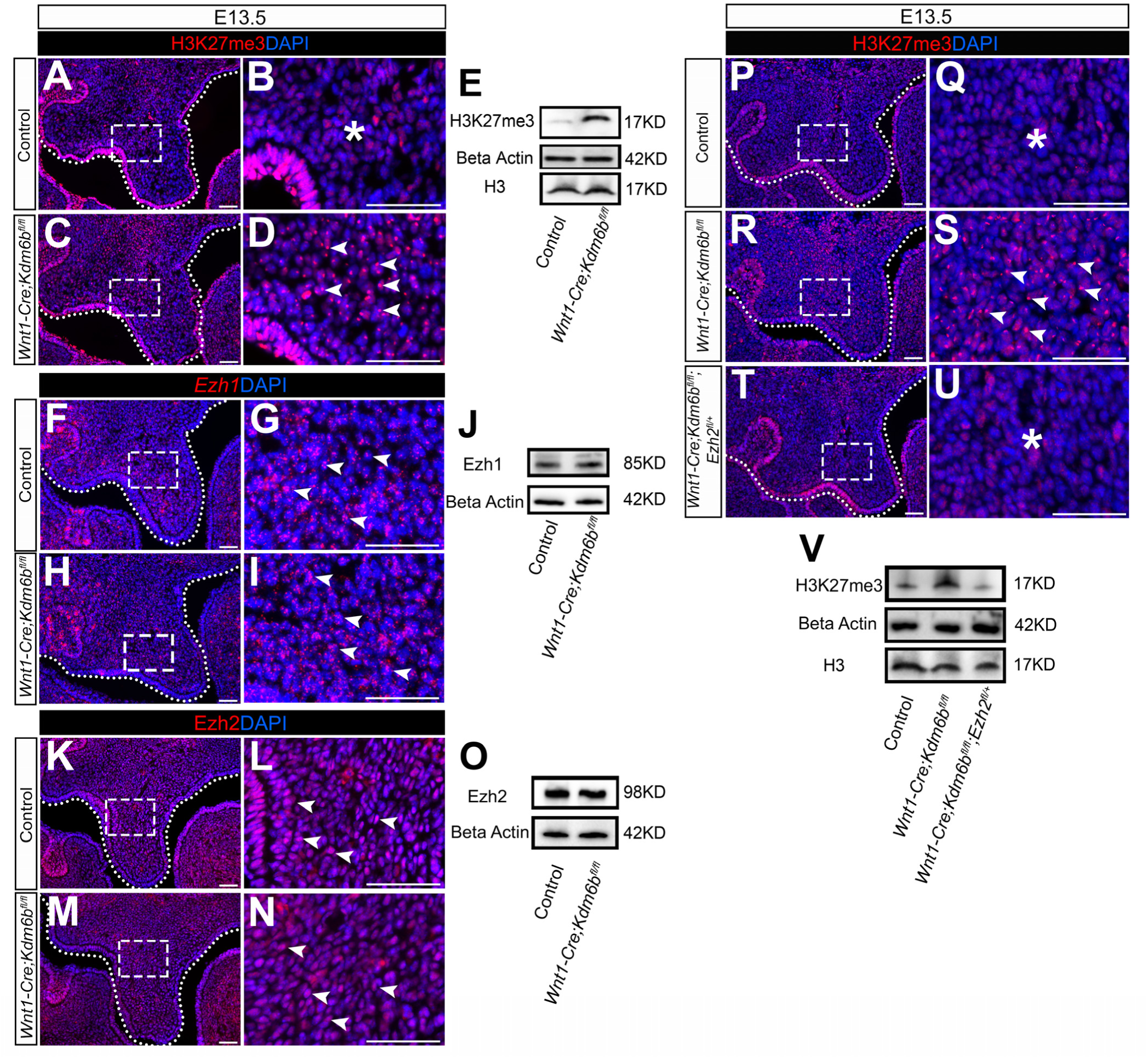
Level of H3K27me3 is precisely regulated by *Kdm6b* and *Ezh2* during palatogenesis. (A-E) Contribution of H3K27me3 in the palatal shelf is evaluated using immunostaining and western blot at E13.5. Dotted lines in A and C indicate palatal shelf region. B and D are magnified images of boxes in A and C. Asterisk in B indicates no accumulation of H3K27me3 observed in control mice. Arrowheads in D indicate accumulation of H3K27me3 observed in *Wnt1-Cre;Kdm6b^fl/fl^* mice. Scale bar: 50 µm. (F-J) Contribution of *Ezh1* in the palatal shelf is evaluated using RNAscope *in situ* hybridization and western blot at E13.5. Dotted lines in F and H indicate palatal shelf region. G and I are magnified images of boxes in F and H. Arrowheads in G and I indicate representative *Ezh1*+ cells. Scale bar: 50 µm.(K-O) Contribution of Ezh2 in the palatal shelf is evaluated using immunostaining and western blot at E13.5. Dotted lines in K and M indicate palatal shelf region. L and N are magnified images of boxes in K and M. Arrowheads in L and N indicate representative Ezh2+ cells. Scale bar: 50 µm. (P-V) Contribution of H3K27me3 in the palatal shelf of control mice, *Kdm6b* mutant mice and Ezh2 haploinsufficient model is evaluated using immunostaining and western blot at E13.5. Dotted lines in P, R, and T indicate palatal shelf region. Q, S, and U are magnified images of boxes in P, R, and T, respectively. Asterisks in Q and U indicate no accumulation of H3K27me3 observed in control (Q) and *Wnt1-Cre;Kdm6b^fl/fl^;Ezh2^fl/+^* mice (U). White arrowheads in S indicate accumulation of H3K27me3 observed in *Wnt1-Cre;Kdm6b^fl/fl^* mice. Scale bar: 50 µm. Figure 5-source data 1 for figure 5E Figure 5-source data 2 for figure 5E Figure 5-source data 3 for figure 5E Figure 5-source data 4 for figure 5E Figure 5-source data 5 for figure 5J Figure 5-source data 6 for figure 5J Figure 5-source data 7 for figure 5J Figure 5-source data 8 for figure 5O Figure 5-source data 9 for figure 5O Figure 5-source data 10 for figure 5O Figure 5-source data 11 for figure 5V Figure 5-source data 12 for figure 5V Figure 5-source data 13 for figure 5V Figure 5-source data 14 for figure 5V

**Figure 6.**
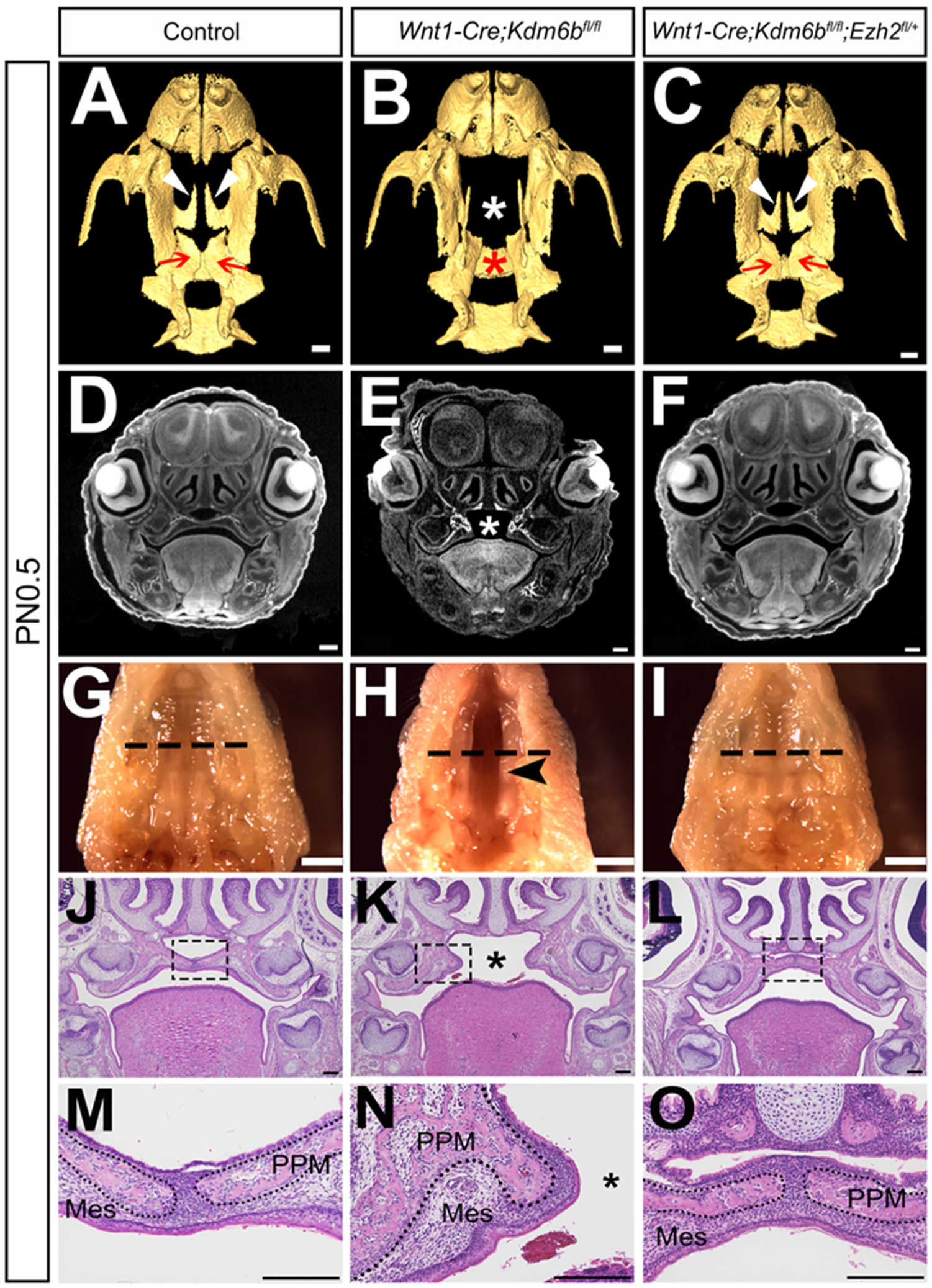
Haploinsufficiency of *Ezh2* in *Wnt1-Cre;Kdm6b^fl/fl^;Ezh2^fl/+^* mice rescues cleft palate. (A-C) CT images at PN0.5. White arrowheads in A and C indicate palatine process of maxilla observed in control and *Wnt1-Cre;Kdm6b^fl/fl^;Ezh2^fl/+^* rescue model. Red arrows in A and C indicate palatine bone observed in control and *Wnt1-Cre;Kdm6b^fl/fl^;Ezh2^fl/+^* rescue model. White asterisk in B indicates missing palate palatine process of maxilla in *Wnt1-Cre;Kdm6b^fl/fl^* mice and red asterisk indicates missing palatine bone in *Kdm6b* mutant mice. Scale bar: 0.4 mm. (D-F) Coronal views of CT images at PN0.5. Asterisk in E indicates cleft palate observed in *Wnt1-Cre;Kdm6b^fl/fl^* mice. Scale bar: 0.3 mm. (G-I) Whole-mount oral view at PN0.5. Arrowhead in H shows complete cleft palate observed in *Wnt1-Cre;Kdm6b^fl/fl^* mice. Dashed lines in G-I indicate location of sections in J-O. Scale bar: 2 mm. (J-O) Histological analysis of samples at PN0.5. Asterisk in K and N indicates cleft palate in *Wnt1-Cre;Kdm6b^fl/fl^* mice. M-O are magnified images of boxes in J-L, respectively. Dotted lines in M-O outline the bone structure. Scale bar: 200 µm. Mes: mesenchyme; PPM: palatine process of maxilla.

### *Kdm6b* activates expression of *p53* through removing H3K27me3 at the promoter of *p53* and providing chromatin accessibility to transcription factor Tfdp1

Chromatin accessibility represents the degree to which chromatinized DNA is able to physically interact with nuclear macromolecules such as transcription factors for gene regulation (Klemm et al. 2019). The repressive mark H3K27me3 is usually associated with facultative heterochromatin and results in transcriptional repression due to decreased chromatin accessibility (Wiles and Selker 2017; Moller et al. 2019; den Broeder et al. 2020). Methyltransferase Ezh2 and demethylase Kdm6a/Kdm6b can both regulate the methylation status of H3K27 to affect gene expression (Pediconi et al. 2019). To test whether *Kdm6b* and *Ezh2* can regulate expression of *p53* via H3K27me3, we first examined whether deposition of H3K27me3 changes at the promoter of *p53* in our models using ChIP-qPCR. A primer set was designed at 1127 bp upstream of *p53* exon1 and the results showed that deposition of H3K27me3 significantly increased at the promoter of *p53* in the palatal region of *Kdm6b* mutant mice, while this increase was dampened in the *Ezh2* haploinsufficiency model (Figure 7A). Meanwhile, haplosufficiency of *Ezh2* in *Wnt1-Cre;Kdm6b^fl/fl^;Ezh2^fl/+^* mice was able to restore the decreased expression of *p53* observed in the CNC-derived palatal mesenchyme of *Wnt1-Cre;Kdm6b^fl/fl^* mice (Figure 7B-H). These data suggested that *Kdm6b* and *Ezh2* co-regulate expression of *p53* through H3K27me3.

**Figure 7.**
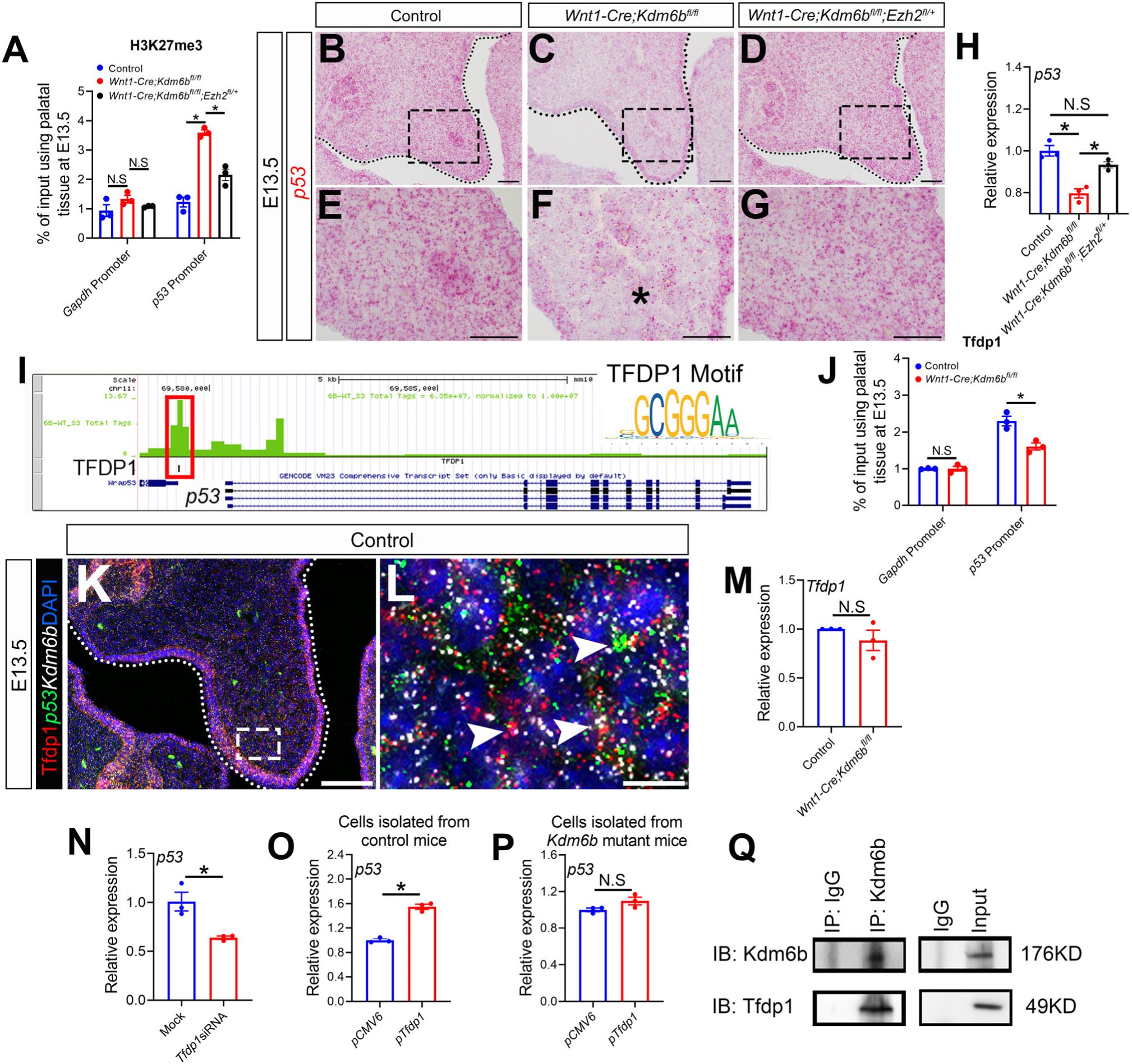
*Kdm6b* regulates expression of *p53* through H3K27me3 and coordinates with transcription factor Tfdp1 in activation of *p53*. (A) ChIP-qPCR shows H3K27me3 deposition at the promoter region of *p53* in palatal tissues of control, *Kdm6b* mutant, and *Ezh2* haploinsufficient mice. ANOVA is used for statistical analysis. Asterisk indicates P < 0.05. (B-G) Expression of *p53* in palatal region at E13.5 using RNAscope *in situ* hybridization. Dotted lines in B, C and D indicate palatal shelf. E, F, and G are magnified images of boxes in B, C, and D, respectively. Asterisk in F indicates decreased expression of p53 observed in *Wnt1-Cre;Kdm6b^fl/fl^* mice. Scale bar: 50 µm. (H) RT-qPCR analysis of *p53* expression in the palatal region of control, *Kdm6b* mutant and *Ezh2* haploinsufficient mice. ANOVA is used for statistical analysis. Asterisk indicates P < 0.05. (I) ATAC-seq analysis indicates that promoter region of *p53* is accessible for transcription factor TFDP1.(J) ChIP-qPCR using palatal tissue shows that binding of Tfdp1 to the promoter of *p53* decreases in the *Kdm6b* mutant mice. (K-L) Co-localization of Tfdp1, *Kdm6b* and *p53* at E13.5 using immunostaining and RNAscope *in situ* hybridization. Dotted lines in K indicate palatal shelf. L is a magnified image of the boxe in K. Arrowheads in L indicate representative cells that are positive for Tfdp1, *Kdm6b* and *p53*. Scale bar: 50 µm in K and 5 µm in L. (M) RT-qPCR quantification shows the expression of *Tfdp1* in samples collected at E13.5. N.S: not significant. (N) RT-qPCR analysis of *p53* expression in palatal mesenchymal cells after *Tfdp1* siRNA transfection. Asterisk indicates P < 0.05. (O-P) RT-qPCR analysis of *p53* expression in palatal mesenchymal cells transfected with *Tfdp1* overexpressing plasmid. Asterisk in O indicates P < 0.05. N.S: not significant. (Q) Co-IP experiment using protein extract from palatal tissues indicates that Kdm6b and Tfdp1 are presented in a same complex. Anti-Kdm6b antibody was used for immunoprecipitation. IgG served as negative control. IP: immunoprecipitation. IB: immunoblotting. Figure 7-source data 1 for figure 7A Figure 7-source data 2 for figure 7H Figure 7-source data 3 for figure 7J Figure 7-source data 4 for figure 7N Figure 7-source data 5 for figure 7O Figure 7-source data 6 for figure 7P Figure 7-source data 7 for figure 7Q Figure 7-source data 8 for figure 7Q Figure 7-source data 9 for figure 7Q Figure 7-source data 10 for figure 7Q Figure 7-source data 11 for figure 7Q

As a H3K27me3 demethylase, Kdm6b is important for the regulation of chromatin structure for gene expression. To target a specific sequence in genome, a histone demethylase needs to interact with DNA binding proteins such as transcription factors or IncRNAs (Dimitrova et al. 2015; Gurrion et al. 2017). To identify a transcription factor that can interact with Kdm6b, we performed ATAC-seq analysis of palate tissue at E13.5. Through motif analysis we found that the promoter region of *p53* was accessible to members of the E2f transcription factor family (E2f4 and E2f6) and transcription factor Tfdp1 (also known as Dp1), a binding partner of E2f family members (Figure 7I and Figure 7-figure supplement 6A). Previous research reported that inactivation of E2fs resulted in milder phenotypes than those associated with loss of *Tfdp1*, which leads to early embryonic lethality (Kohn et al. 2003). This result suggested that Tfdp1 may play a more critical role than E2fs during embryonic development. A motif of Tfdp1 was detected 1011 bp upstream of *p53* exon 1, which is very close to the H3K27me3 deposition site, by ATAC-seq analysis. ChIP-qPCR using palate tissue at E13.5 also revealed that binding of Tfdp1 to the promoter region of *p53* decreased in the *Kdm6b* mutant mice (Figure 7J). Immunohistochemistry analysis showed that Tfdp1+ cells were distributed in the palatal region and co-expressed with *p53* and *Kdm6b* (Figure 7K-L). We also confirmed co-expression of *p53* and *Tfdp1* in the palatal region using our previously published scRNA-seq data (Figure 7-figure supplement 6B) (Han et al. 2021). Meanwhile, expression level and distribution of Tfdp1 was not affected in the palatal mesenchyme in *Wnt1-Cre;Kdm6b^fl/fl^* mice (Figure 7M and Figure 7-figure supplement 6C-D). These data indicated that *Tfdp1* is not a downstream target of *Kdm6b*. To further test whether *Tfdp1* regulated expression of *p53* in the palatal mesenchymal cells, we transfected palatal mesenchymal cells from control mice at E13.5 using siRNA to knock down *Tfdp1*. qPCR revealed that the expression of *p53* was decreased in cells treated with siRNA for *Tfdp1* (Figure 7-figure supplement 6E and Figure 7N). This data further indicated that *p53* is a direct downstream target of *Tfdp1*.

To reveal the function of *Kdm6b-Tfdp1* interaction in the regulation of *p53* during palatogenesis, we transfected palatal mesenchymal cells with *Tfdp1*-overexpressing plasmid and found that expression of *p53* increased in the cells from control mice but not in the cells from *Wnt1-Cre;Kdm6b^fl/fl^* mice (Figure 7-figure supplement 6F-G and Figure 7O-P). This result suggested that *Kdm6b* plays an essential role in activation of *p53* through interaction with *Tfdp1* during palatogenesis. To further test whether Kdm6b and Tfdp1 are present in a same complex, we performed Co-IP experiments and found that these two proteins were indeed involved in the same complex (Figure 7Q). Collectively, these data suggested that Kdm6b and Tfdp1 work together to activate *p53* expression in the palatal mesenchyme and play an important role in regulating palatogenesis (Figure 8).

**Figure 8.**
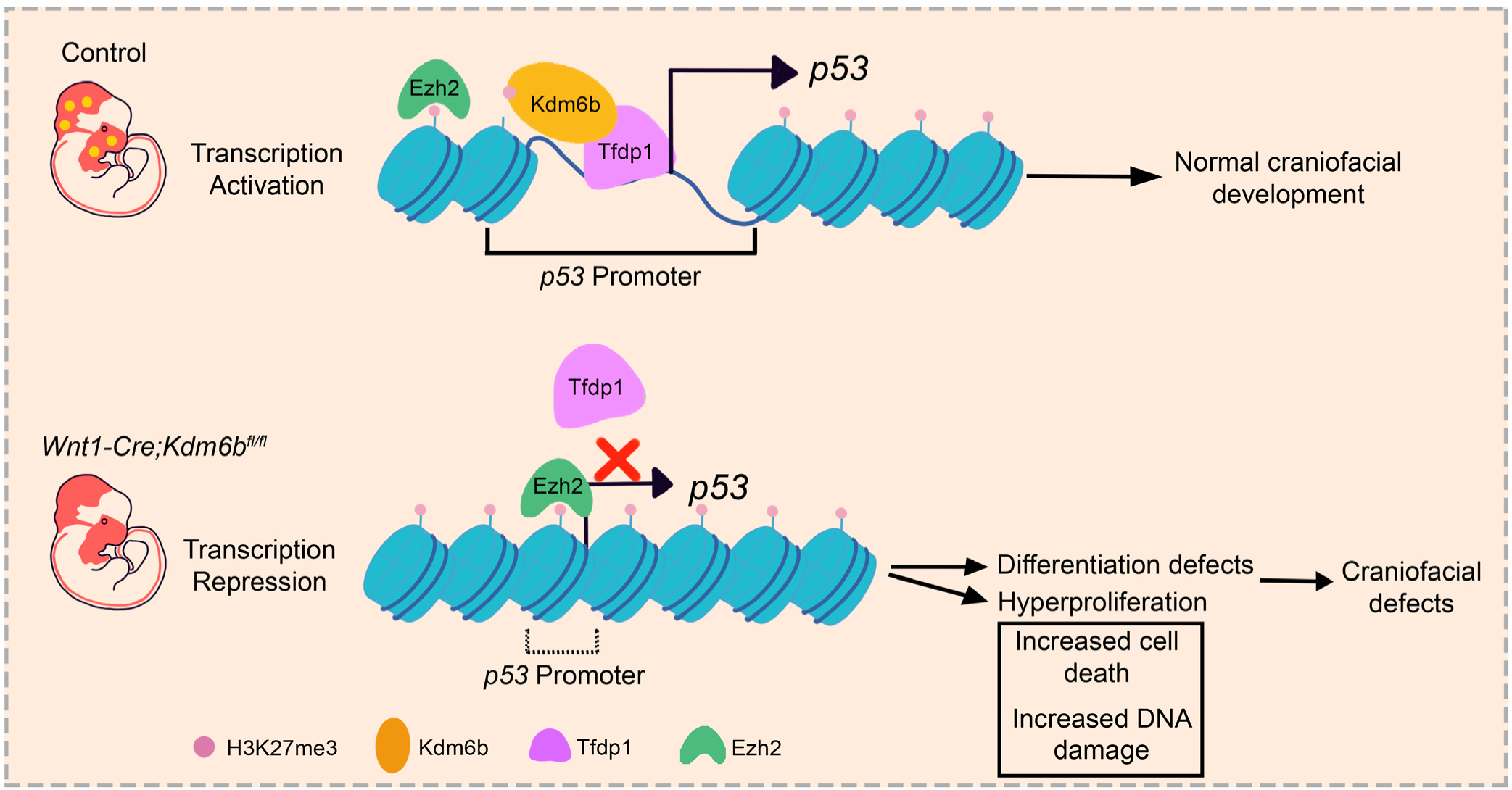
Summary schematic drawing.

## Discussion

The development of an organism from a single cell to multiple different cell types requires tightly regulated gene expression (Bruneau et al. 2019). Transcription factors, which are among the key regulators of this process, are intimately involved in cell fate commitment (Nelms and Labosky 2010; Soldatov et al. 2019). However, a transcription factor by itself cannot act on densely packed DNA in chromatin form. Thus, transcription factors must work in coordination with epigenetic regulatory mechanisms such as histone modifications, DNA methylation, chromatin remodeling and others to dynamically regulate chromatin states for gene expression (Wilson and Filipp 2018; Gokbuget and Blelloch 2019). Insults to the epigenetic landscape due to genetic, environmental or metabolic factors can lead to diverse developmental defects and diseases (Hobbs et al. 2014; Zoghbi and Beaudet 2016; Flavahan et al. 2017). Cleft palate comprises 30% of orofacial clefts, and can result from genetic mutations, environmental effects, or a combination thereof (Seelan et al. 2012). Much progress has been made in taking inventory of the gene mutations associated with craniofacial defects in recent years, and growing evidence has shown that epigenetic regulation plays an important role during neural crest development. For example, haploinsufficiency of *KDM6A* in humans causes severe psychomotor developmental delay, global growth restriction, seizures and cleft palate (Lindgren et al. 2013). Furthermore, studies have shown that *Kdm6a* and *Arid1a* are both indispensable during neural crest development (Chandler and Magnuson 2016; Shpargel et al. 2017). DNA methyltransferase3A (DNMT3A) plays a critical role in mediating the transition from neural tube to neural crest fate (Hu et al. 2012). Meanwhile, loss of *Ezh2*, a component of PRC2, in CNC-derived cells completely prevents craniofacial bone and cartilage formation (Schwarz et al. 2014). These studies have clearly shown that epigenetic regulation is crucial for neural crest development. In this study, we further demonstrate the important role of epigenetic regulation during the neural crest contribution to palate development using *Wnt1-Cre;Kdm6b^fl/fl^* mice as a model. We show that the demethylase Kdm6b is not only required for normal CNC-derived palatal mesenchymal cell proliferation, but also for maintaining cell differentiation.

Epigenetic regulators, transcription factors, and lineage-specific genes work together to achieve spatiotemporally restricted, tissue-specific regulation (Hu et al. 2014). In this study, we reveal that *Kdm6b* works with the transcription factor *Tfdp1* to specifically regulate the expression of *p53.* The molecular mechanisms underlying the function of *p53* in genomic stability and tumor suppression have been studied extensively. However, the role of *p53* in regulating the development of CNC-derived cells still remains largely unclear, although several studies have been conducted recently on certain aspects of this topic. For instance, it has been shown that *p53* is able to coordinate CNC cell growth and epithelial-mesenchymal transition/delamination processes by modulating cell cycle genes and proliferation(Rinon et al. 2011). It has also been established that both deletion and overexpression of *p53* result in craniofacial defects (Tateossian et al. 2015; Bowen et al. 2019). Furthermore, nuclear stabilization of p53 protein in *Tcof^+/-^* mice induces neural crest cell progenitors to undergo cell-cycle arrest and caspase3-mediated apoptosis in the neuroepithelium. Inhibition of p53 function successfully rescues the neurocristopathy in an animal model of Treacher Collins syndrome, which results from mutation in *Tcof1* (Jones et al. 2008). These studies have clearly shown that appropriate function of *p53* is indispensable in CNCCs. However, none of these studies have addressed upstream regulation of *p53* in CNCCs.

Here, we show that proper function of *p53* during the differentiation and proliferation of CNCCs is orchestrated by *Kdm6b* and *Ezh2* through H3K27me3. Altering the balance between *Ezh2* and *Kdm6b* can cause abnormal H3K27me3 function, which further affects the downstream transcription factor *p53*. In addition, we have detected spontaneous DNA damage in the developing palate and increased accumulation of DNA damage in the *Wnt1-Cre;Kdm6b^fl/fl^* mice. These findings further demonstrate the critical function of *p53* in protecting embryonic cells from DNA damage during development.

Furthermore, the ability of cells to proliferate is limited by the length of the telomeres, which gradually shorten during each cell replication (Blagoev 2009). Once the telomeres are too short for DNA replication, the result is cellular senescence, which induces an irreversible inability to proliferate (Bernadotte et al. 2016). In this study we notice that downregulated expression of *p53* in *Wnt1-Cre;Kdm6b^fl/fl^* mice results in hyperproliferation and increased DNA damage in the proliferative cells, which might further leads to cell senescence.

Previous research has shown that KDM3A functions as a cofactor of STAT3 to activate the JAK2-STAT3 signaling pathway (Kim et al. 2018) and that KDM2A coordinates with c-Fos in regulating *COX-2* (Lu et al. 2015). Our study shows that Kdm6b coordinates with the transcription factor Tfdp1 to activate expression of *p53* in CNCCs, and that *Ezh2* and *Kdm6b* co-regulate H3K27 methylation status, which may affect the ability of Tfdp1 to bind to the chromatin during palatogenesis. It has been reported that *Tfdp1* is crucial for embryonic development and for regulating Wnt/β-catenin signaling (Kohn et al. 2003; Kim et al. 2012). Interaction between Kdm6b and Tfdp1 discovered in this study further increases our knowledge of the coordination between epigenetic regulators and transcription factors during organogenesis. As environmental insults can adversely affect the function of epigenetic regulators, our findings provide a better understanding of the epigenetic regulation and transcription factors involved in regulating the fate of CNC cells and craniofacial development, which can provide important clues about human development, as well as potential therapeutic approaches for craniofacial birth defects.

## Materials and Methods

### Animals

To generate *Wnt1-Cre;Kdm6b^fl/fl^* mice, we crossed *Wnt1-Cre;Kdm6b^fl/+^* mice with *Kdm6b^fl/fl^* mice (Zhao et al. 2008; Manna et al. 2015). Reporter mice used in this study were tdTomato conditional reporter (JAX#007905) (Madisen et al. 2010). *Ezh2^fl/fl^* and *p53^fl/fl^* mice were purchased from Jackson Laboratory (JAX#022616, #008462) (Marino et al. 2000; Shen et al. 2008). Genotyping was carried out as previously described (Zhao et al. 2008). Briefly, tail samples were lysed by using DirectPCR tail solution (Viagen 102 T) with overnight incubation at 55°C. After heat inactivation at 85°C for 1 hour, PCR-based genotyping (GoTaq Green MasterMix, Promega, and C1000 Touch Cycler, Bio-rad) was used to detect the genes. All mouse studies were conducted with protocols approved by the Department of Animal Resources and the Institutional Animal Care and Use Committee (IACUC) of the University of Southern California (Protocols 9320 and 20299).

### MicroCT analysis

MicroCT was used to analyze the control, *Kdm6b,* and other mutant samples. Mouse samples were dissected and fixed in 4% paraformaldehyde overnight at 4°C followed by CT scanning (Scanco Medical µCT50 scanner) at the University of Southern California Molecular Imaging Center as previously described (Grosshans et al. 2006; Sugii et al. 2017). AVIZO 9.1.0 (Visualization Sciences Group) was used for visualization and 3D microCT reconstruction.

### Alcian blue-Alizarin red staining

Mouse heads were dissected and fixed in 95% EtOH overnight at room temperature. Staining was performed as previously described (Rigueur and Lyons 2014). Briefly, 95% EtOH was replaced with 100% acetone for 2 days and then samples were incubated in Alcian blue solution (80% EtOH, 20% glacial acetic acid, and 0.03% (w/v) Alcian blue 8GX (Sigma, A3157) for 1-3 days. Samples were then destained with 70% EtOH and incubated in 95% EtOH overnight. After incubation, samples were pre-cleared with 1% KOH and then incubated in Alizarin red solution (0.005% (w/v) Alizarin red (Sigma, A5533) in 1% (w/v) KOH) for 2-5 days. After clearing samples with 1% KOH, they were stored in 100% glycerol until analysis.

### Sample preparation for sectioning

Samples for paraffin sectioning were prepared using the standard protocol in our laboratory. Briefly, samples were fixed in 4% PFA and decalcified with 10% EDTA as needed. Then, samples were dehydrated with serial ethanol solutions (50%, 70%, 80%, 90% and 100%) at room temperature followed by xylene and then embedded in paraffin wax. Sections were cut to 6 µm on a microtome (Leica) and mounted on SuperFrost Plus slides (Fisher, 48311-703). Cryosectioning samples were fixed and decalcified the same way as samples prepared for paraffin sectioning. Sucrose (15% and 30%) was used to remove water from the samples before embedding them in OCT compound (Tissue-Tek, 4583). Cryosections were cut to 8 µm on a cryostat (Leica) and mounted on SuperFrost Plus slides (Fisher).

### Histological analysis

Paraffin sections prepared as described above were used for histological analysis. Hematoxylin and Eosin staining were performed using the standard protocol (Cardiff et al. 2014).

### Immunofluorescence assay

Cryosections and paraffin sections prepared as described above were used for immunofluorescence assays. Sections were dried for 2 hours at 55°C. Paraffin sections were deparaffinized and rehydrated before antigen retrieval. Heat mediated antigen retrieval was used to process sections (Vector, H-3300) and then samples were blocked for 1 hour in blocking buffer at room temperature (PerkinElmer, FP1020). Primary antibodies diluted in blocking buffer were incubated with samples overnight at 4°C. After washing with PBST (0.1% Tween20 in 1xPBS), samples were then incubated with secondary antibodies at room temperature for 2 hours. DAPI (Sigma, D9542) was used for nuclear staining. All images were acquired using Leica DMI 3000B and Keyence BZ-X710/810 microscopes. Detailed information about primary and secondary antibodies is listed in Table S1.

### EdU labeling

EdU solution was prepared at 10 mg/mL in PBS, and then pregnant mice at the desired stage were given an intra-peritoneal injection (IP) based on their weight (0.1 mg of EdU/1 g of mouse). Embryos were collected after 2 hours or 48 hours and then prepared for sectioning as above. EdU signal was detected using Click-It EdU cell proliferation kit (Invitrogen, C10337) and images were acquired using Leica DMI 3000B and Keyence BZ-X710/810 microscopes.

### TUNEL staining

Cryosections and paraffin sections were prepared as described above, and cell death was analyzed using TUNEL staining according to the manufacturer’s protocol (Invitrogen, C10245). Images were acquired using Keyence BZ-X710/810 microscope.

### RNAscope *in situ* hybridization

RNAscope *in situ* hybridization in this study was performed on cryosections using RNAscope 2.5HD Reagent Kit-RED assay (Advanced Cell Diagnostics, 322350) and RNAscope multiplex fluorescent v2 assay (Advanced Cell Diagnostics, 323100) according to the manufacturer’s protocol. RNAscope probes used in this study included *Kdm6a*, *Kdm6b, Uty,* and *p53*. Detailed information about probes is listed in Table S2.

### RNA-sequencing and analysis

Palate samples from control and *Wnt1-Cre;Kdm6b^fl/fl^* mice were collected at E12.5 for RNA isolation with RNeasy Micro Kit (Qiagen) according to the manufacturer’s protocol. The quality of RNA samples was determined using an Agilent 2100 Bioanalyzer and all samples for sequencing had RNA integrity (RIN) numbers > 9. cDNA library preparation and sequencing were performed at the USC Molecular Genomics Core. Single-end reads with 75 cycles were performed on Illumina Hiseq 4000 equipment and raw reads were trimmed and aligned using TopHat (Version 2.0.8) with the mm10 genome. CPM was used to normalize the data and differential expression was calculated by selecting transcripts that changed with p < 0.05.

### RNA extraction and real-time qPCR

Palatal tissue used for RNA isolation was dissected at desired stages and an RNeasy Plus Micro Kit (Qiagen, 74034) was used to isolate the total RNA followed by cDNA synthesis using an iScript cDNA synthesis kit (Bio-Rad, 1708891). Real-time qPCR quantification was done on a Bio-Rad CFX96 Real-Time system using SsoFast EvaGreen Supermix (Bio-Rad, 1725201). Detailed information on primers is listed in Table S3.

### ChIP-qPCR

Palate tissue was dissected from control and *Wnt1-Cre;Kdm6b^fl/fl^* mice at E13.5. Each replicate contained 60-80 mg tissue combined from multiple animals. Samples were prepared following the manufacturer’s protocol (Chromatrap, 500191). Briefly, tissue was cut into small pieces and then fixed with 1% formaldehyde at room temperature for 15 minutes, followed by incubating with 0.65 M glycine solution. Then the sample was washed twice with PBS, resuspended in Hypotonic Buffer and incubated at 4°C for 10 minutes to obtain nuclei, which were then resuspended in Digestion Buffer. After chromatin was sheared to 100-500 bp fragments using Shearing Cocktail, 10 µg chromatin with H3K27me3 antibody (CST 9733s, 1:50), DP1 antibody (Abcam ab124678, 1:10) or Immunoglobulin G negative control (2 µg) was added to Column Conditioning Buffer to make up the final volume of 1000 µl. Immunoprecipitation (IP) slurry was mixed thoroughly and incubated on an end-to-end rotor for 1 hour at 4 °C. An equivalent amount of chromatin was set as an input. After 1 hour incubation, IP slurry was purified using Chromatrap® spin column at room temperature and chromatin was eluted using ChIP-seq elution buffer. Chromatin sample and input were further incubated at 65 °C overnight to reverse cross-linking. DNA was purified with Chromatrap® DNA purification column after proteinase K treatment. ChIP eluates, negative control and input were assayed using real-time qPCR. Primers were designed using the promoter region of *p53*. Detailed information is available in Table S3.

### Western blot and Co-Immunoprecipitation

For western blot, palate tissue was dissected from control and *Wnt1-Cre;Kdm6b^fl/fl^* mice at E13.5. The tissue sample was lysed using RIPA buffer (Cell Signaling, 9806) with protease inhibitor (Thermo Fisher Scientific, A32929) for 20 minutes on ice followed by centrifugation at 4°C to remove tissue debris. Protein extracts were then mixed with sample buffer (Bio-Rad, 1610747) and boiled at 98°C for 10 minutes. Then denatured protein extract was separated in 4%-15% precast polyacrylamide gel (Bio-Rad, 456-1084) and then transferred to 0.45µm PVDF membrane. Transferred membrane was incubated with 5% milk for 1 hour at room temperature, and incubated with primary antibody (Table S4) at 4°C overnight. After washing with TBST, membrane was incubated with secondary antibody for 2 hours at room temperature and signals were detected using SuperSignal West Femto (Thermo Fisher Scientific, 34094) and Azure 300 (Azure Biosystems).

For CoIP, palate tissue was dissected from control mice at E13.5 and 60-80 mg tissue was combined as one sample for each replicate. After lysing using RIPA buffer, 60 µl of the protein extract was mixed with sample buffer and boiled at 98 °C to serve as input. The remaining protein extract was incubated with primary antibody at 4 °C overnight. Protein G beads from GE Healthcare (GE Healthcare, 10280243) were used to purify the target protein and then the protein sample was analyzed using western blot. Detailed information about primary and secondary antibodies is listed in Table S4.

### siRNA and plasmid transfection

Palatal tissue was dissected from control and *Wnt1-Cre;Kdm6b^fl/fl^* mice at E13.5, then cut into small pieces using a scalpel. This minced tissue was then cultured in DMEM medium (Gibco, 2192449) containing 40% MSC FBS (Gibco, 2226685P) and 1% Pen Strep (Gibco, 2145477) at 37°C.

siRNA (Qiagen) and plasmid (OriGene) transfection was performed following the manufacturer’s protocol (Qiagen, 301704 and OriGene, TF81001). Briefly, siRNA was transfected into cells in 24-well plates at 10nM for 3 days followed by qPCR and EdU proliferation assay. Plasmid was transfected into cells in 24-well plates using 1µg/µl stock solution for 2 days followed by real-time qPCR. Primers designed for qPCR are listed in Table S3. siRNA sequence and plasmid information are listed in Table S5 and Table S6.

### ATAC-seq analysis

Palate tissue of E13.5 control mice was digested using TrypLE express enzyme (Thermo Fisher Scientific, 12605010) and incubated at 37 °C for 20 minutes with shaking at 600 rpm. Single-cell suspension was prepared according to the 10X Genomics sample preparation protocol and processed to generate ATAC-seq libraries according to a published protocol (Buenrostro et al. 2015). Sequencing was performed using the NextSeq 500 platform (Illumina) and ATAC-seq reads were aligned to the UCSC mm10 reference genome using BWA-MEN (Li 2013). Then ATAC-seq peaks were called by MACS2 and annotated. Known transcription factor biding motifs were analyzed by HOMER (Zhang et al. 2008; Heinz et al. 2010). Quality files for sequencing is listed in Table S7

### Cell differentiation assay

Palatal tissue was dissected from control and *Wnt1-Cre;Kdm6b^fl/fl^* mice at E13.5 and cultured as previously described. Then the differentiation assay was conducted according to the manufacturer’s protocol (Gibco, A1007201). Briefly, mesenchymal cells were seeded into cell culture plates at the desired concentration followed by incubation at 36°C in a humidified atmosphere of 5% CO_2_ for the required time (a minimum of 2 hours and up to 4 days). Then growth medium was replaced by complete differentiation medium and cells were continuously incubated for 3 weeks under osteogenic conditions. After specific periods of cultivation, cells were stained using 2% Alizarin Red S solution (PH 4.2) solution. Images were acquired using EPSON Scan and Keyence BZ-X710/810 microscopes. Quantification of the Alizarin Red S staining was conducted according to the manufacturer’s protocol (ScienCell, 8678).

### Nutlin-3 treatment

Nutlin-3 (Sigma, N6287) was dissolved in corn oil (Sigma, C8267) with 10% DMSO (Sigma, D2650) and given to pregnant mice on days 10.5, 12.5 and 14.5 of pregnancy at a dosage based on their weight (10 mg/kg) (Li et al. 2016). Then embryos were collected at E16.5 for analysis.

### Statistics

Statistical analysis was completed using GraphPad Prism and significance was assessed by independent two-tailed Student’s t-test or ANOVA. The chosen level of significance for all statistical tests in this study was p <0.05. Data is presented as mean ± SEM. N=3 samples were analyzed for each experimental group unless otherwise stated.

## Acknowledgements

We thank Bridget Samuels and Linda Hattemer for critical reading of the manuscript and also acknowledge USC Libraries Bioinformatics Service for their assisting with data analysis. Meanwhile, we thank USC Office of Research and the Norris Medical Library for the bioinformatics software and computing resources. Furthermore, we appreciate the funding support from the National Institute of Dental and Craniofacial Research, National Institutes of Health (R01 DE012711, R01 DE022503, and U01 DE028729 to Yang Chai).

## Author contributions

T.G. and Y.C. designed the study. T.G. carried out most of the experiments, generated all figures and analyzed the data. X.H., J.H., J.F., and J.J. participated in the sample collection and data analysis. E.J., and J.L. provided critical comments for this manuscript. T-V.H. participated in the microCT analysis. J.X. provided critical suggestion for the manuscript. T.G. and Y.C. co-wrote the paper. Y.C. supervised the research.

## Supplementary figures

**Figure 1-figure supplement 1.**
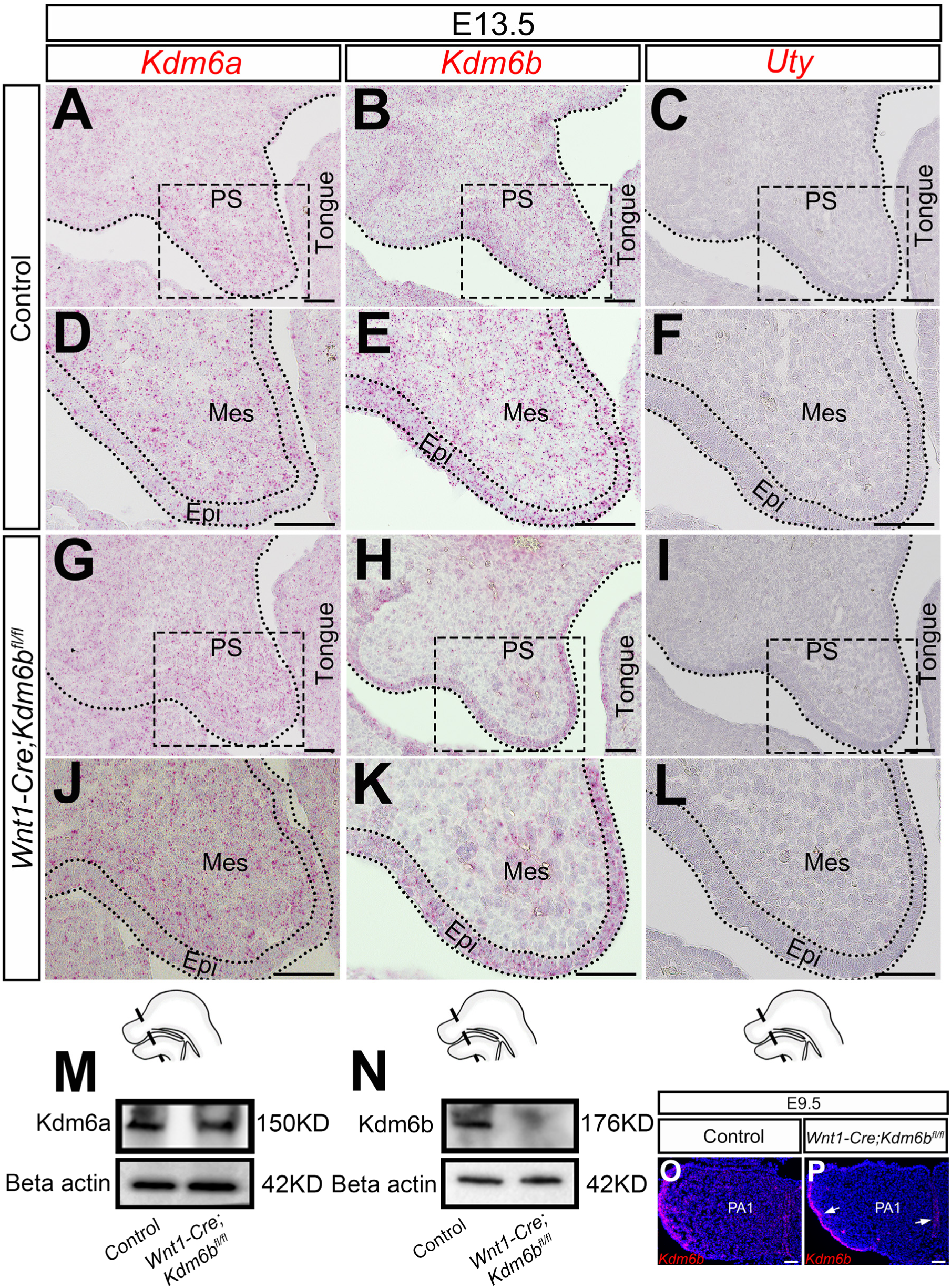
Expression of Kdm6 family. (A-L) Expression of Kdm6 family genes in the palatal region at E13.5 using RNAscope *in situ* hybridization. D, E, F, J, K, and L are magnified images of boxes in A, B, C, G, H, and I respectively. Dotted lines in A, B, C, G, H, and I indicate region of palatal shelf. Dotted lines in D, E, F, J, K, and L indicate epithelium. Schematic drawing at bottom of the figure indicates the location of the presented section. Scale bar: 50 µm. PS: palatal shelf; Mes: mesenchyme; Epi: epithelium. (M-N) Protein quantification of Kdm6a and Kdm6b in control and *Kdm6b* mutant palatal region at E13.5 using western blot. (O-P) Expression of *Kdm6b* in the first pharyngeal arch at E9.5 assessed using RNAscope *in situ* hybridization. Arrows in P indicate expression of *Kdm6b* at epithelium. PA1: first pharyngeal arch. Scale bar: 50 µm. Figure supplement 1-source data 1 for figure supplement 1M Figure supplement 1-source data 2 for figure supplement 1M Figure supplement 1-source data 3 for figure supplement 1M Figure supplement 1-source data 4 for figure supplement 1N Figure supplement 1-source data 5 for figure supplement 1N Figure supplement 1-source data 6 for figure supplement 1N

**Figure 1-figure supplement 2.**
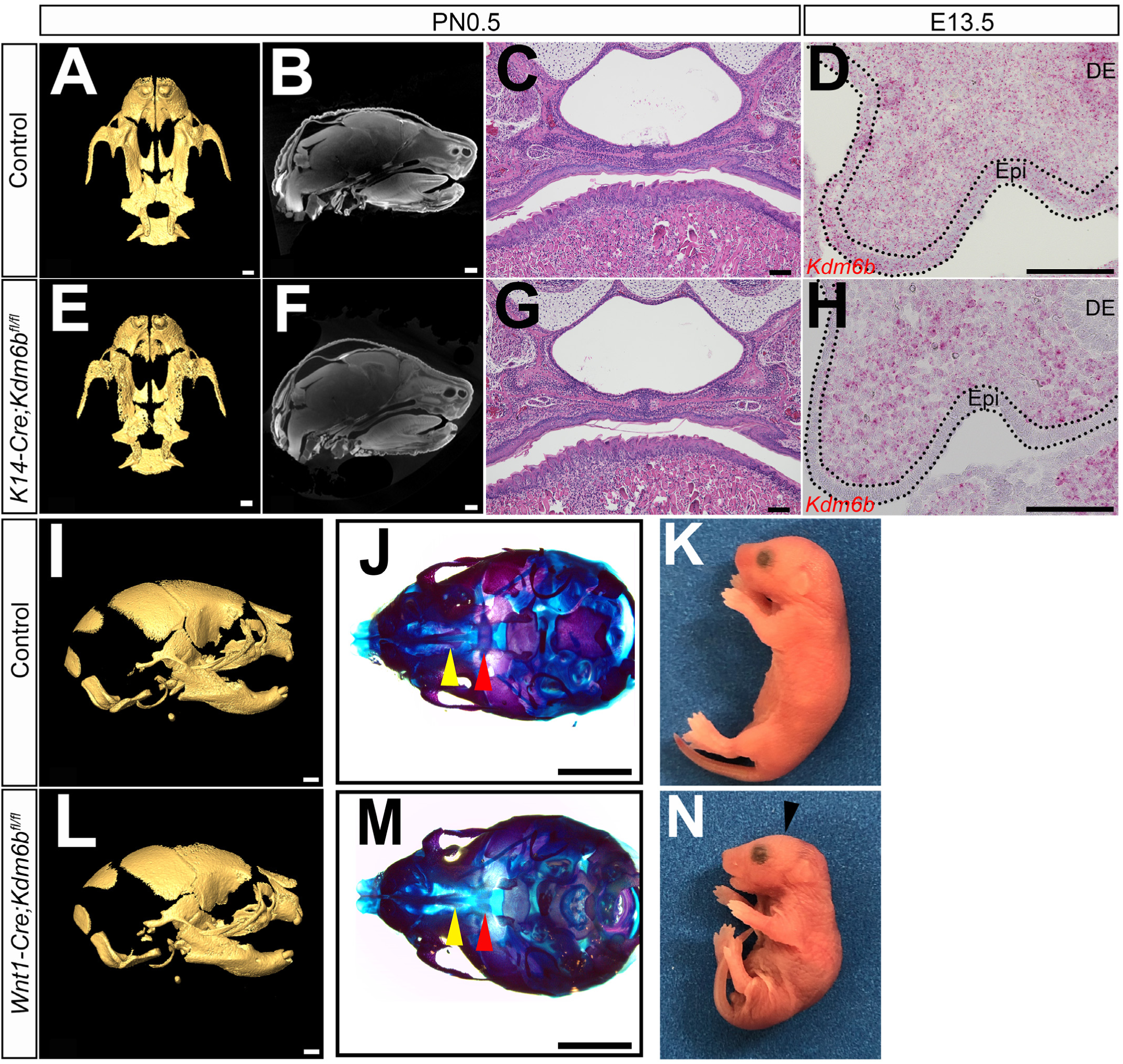
Loss of *Kdm6b* in epithelium and CNC-derived cells. (A-C and E-G) CT images and histological analysis of *K14-Cre;Kdm6b^fl/fl^* mice at PN0.5. Scale bars in A and E: 0.4 mm; Scale bars in B and F: 0.6 mm; Scale bars in C and G: 100 µm. (D and H) Expression of *Kdm6b* assessed using RNAscope *in situ* hybridization at E13.5. *Kdm6b* is efficiently knocked out from epithelium in *K14-Cre;Kdm6b^fl/fl^* mice. Dotted lines in D and H indicate epithelium. Scale bar: 100 µm. Epi: epithelium; DE: dental epithelium. (I-N) No obvious phenotype was observed in skull bones or mandible in *Wnt1-Cre;Kdm6b^fl/fl^* mice at PN0.5. Yellow triangles in J and M indicate location of palatine process of maxilla and red triangle indicates location of palatine bone. Arrowhead in N indicates flatten skull observed in *Wnt1-Cre;Kdm6b^fl/fl^* mice. Scale bar in I and L: 0.6 mm; Scale bar in J and M: 1 mm.

**Figure 1-figure supplement 3.**
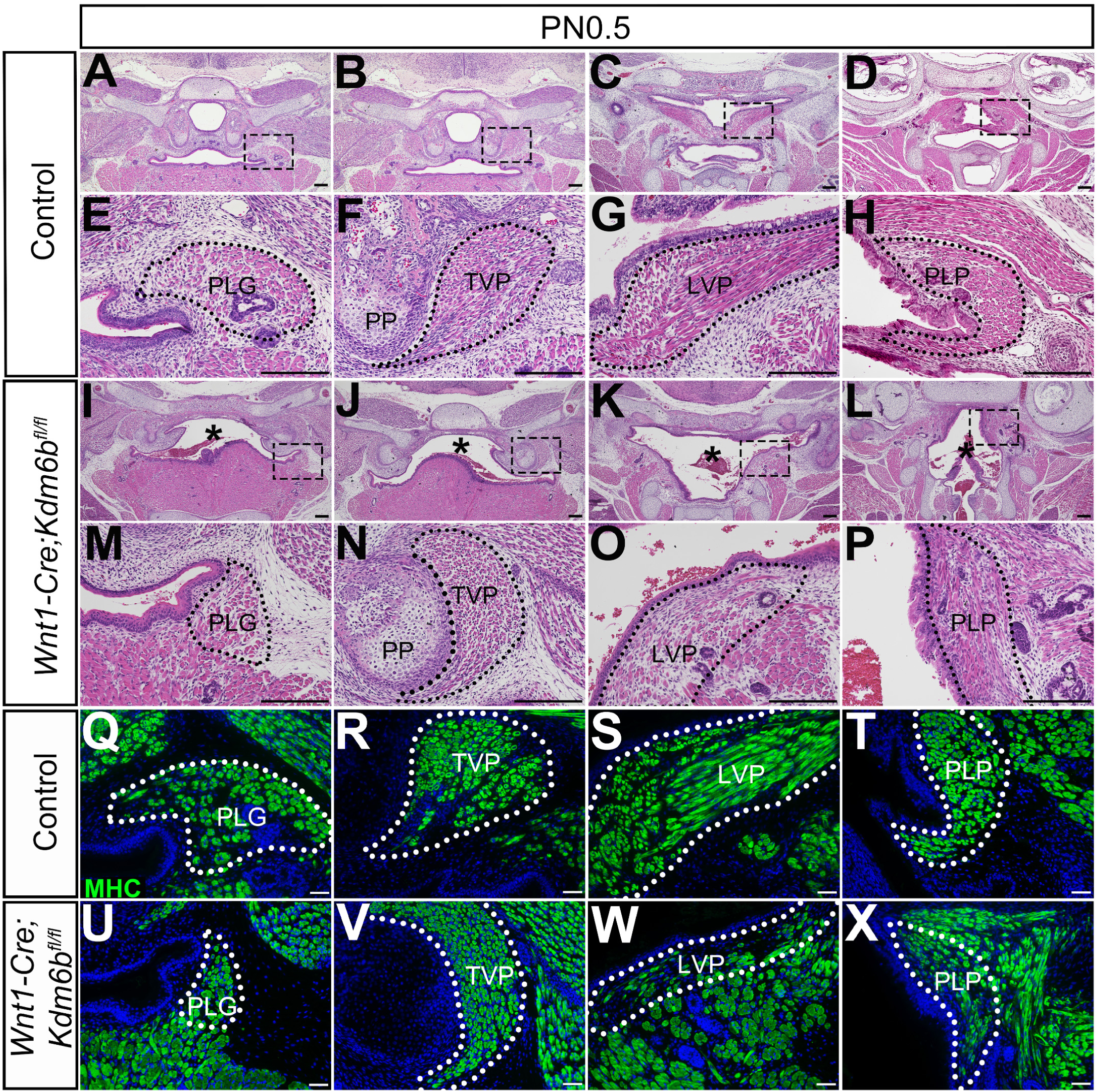
Loss of *Kdm6b* in CNC-derived cells results in soft palate muscle defects. (A-P) Histological analysis of soft palate muscles at PN0.5. Boxes in A, B, C, D, I, J, K, and L are shown magnified in E, F, G, H, M, N, O, and P, respectively. Dotted lines outline each muscle. Asterisks in I, J, K, and L indicate cleft palate observed in *Wnt1-Cre;Kdm6b^fl/fl^* mice. PP: pterygoid plate; PLG: palatoglossus; TVP: tensor veli palatini; LVP: levator veli palatini; PLP: palatopharyngeus. Scale bar: 100 µm. Asterisks in I-L indicate cleft in *Wnt1-Cre;Kdm6b^fl/fl^* mice. (Q-X) Immunostaining of MHC at PN0.5. Dotted lines outline each muscle. PLG: palatoglossus; TVP: tensor veli palatini; LVP: levator veli palatini; PLP: palatopharyngeus. Scale bar: 50 µm.

**Figure 2-figure supplement 4.**
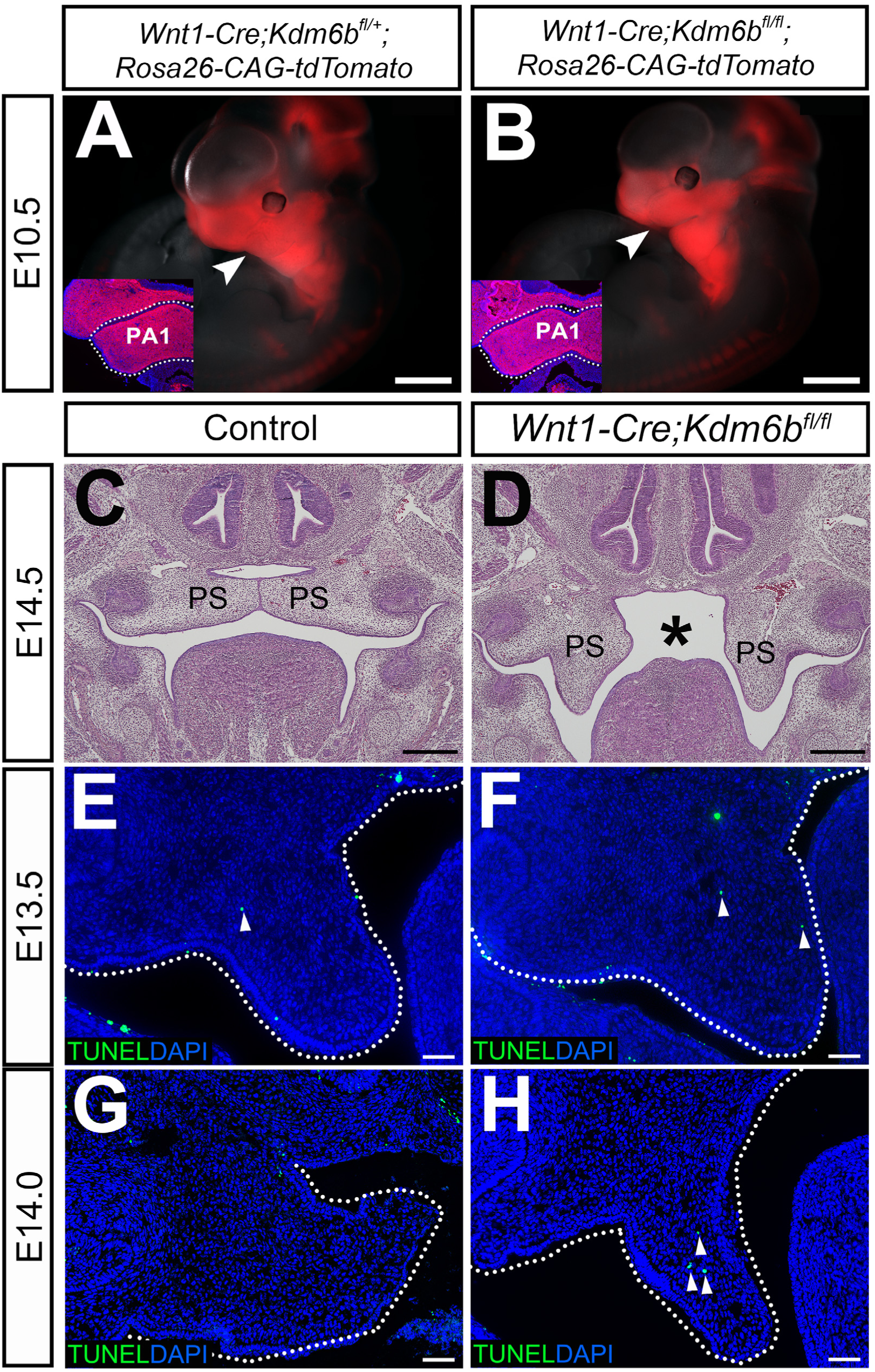
*Kdm6b* is not required for CNCCs to populate pharyngeal arches but is critical for survival of palatal mesenchymal cells. (A-B) Whole-mount images of tdTomato reporter mice at E10.5. Arrowheads indicate CNCCs that have successfully migrated to the pharyngeal arch at E10.5. No differences were observed between control and *Wnt1-Cre;Kdm6b^fl/fl^* mutant mice. Insets show immunostaining of tdTomato at E10.5. Dotted lines in the insets indicate first pharyngeal arch. PA1: first pharyngeal arch. Scale bars: 1 mm. (C-D) Histological analysis of samples at E14.5. Asterisk in D indicates cleft palate observed in *Wnt1-Cre;Kdm6b^fl/fl^* mice. PS: palatal shelf. Scale bar: 100 µm. (E-H) TUNEL staining at E13.5 and E14. Dotted lines indicate palatal shelves. Arrows in E, F and H indicate cells with positive TUNEL staining. Scale bar: 50 µm.

**Figure 4-figure supplement 5.**
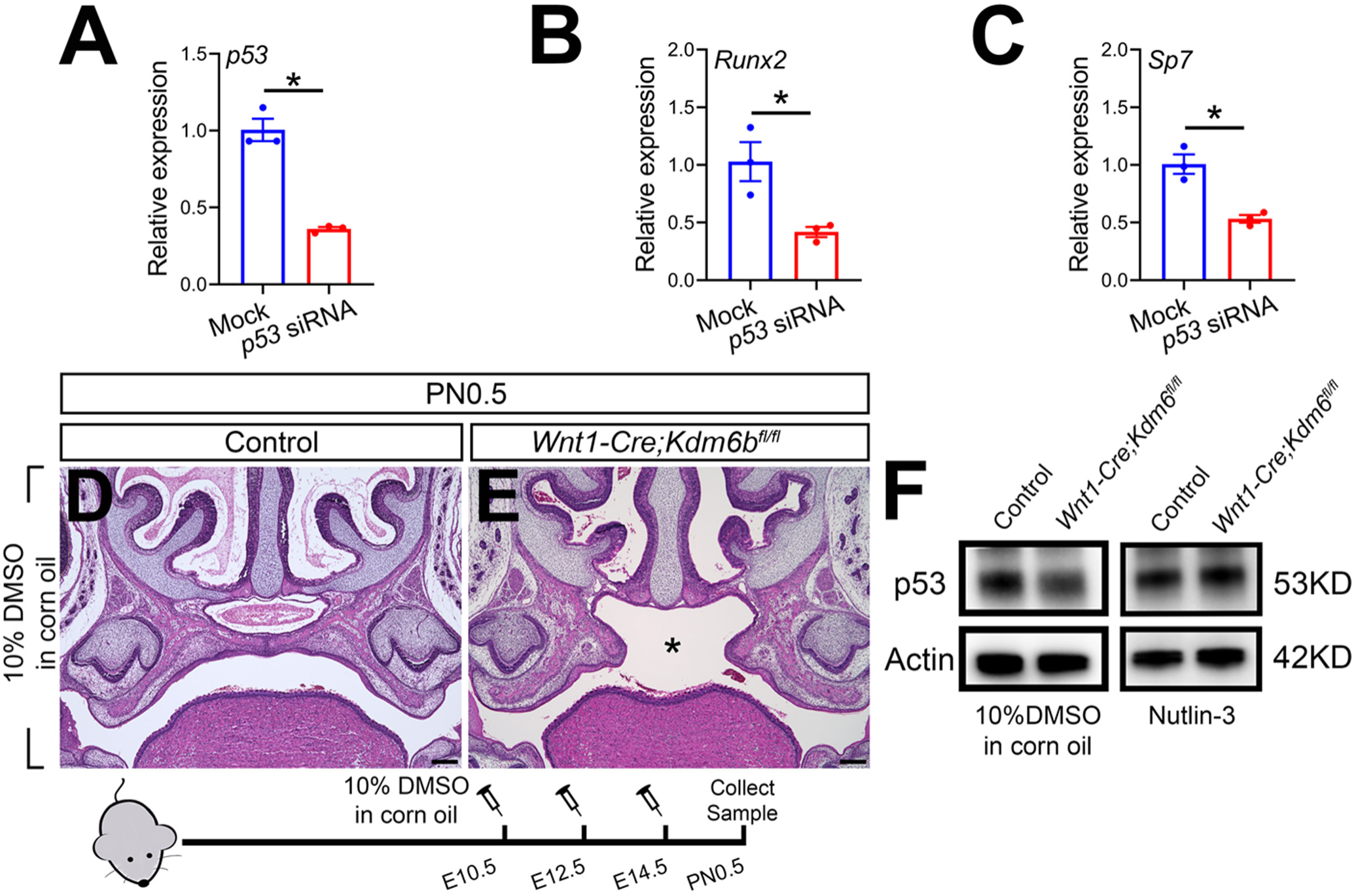
*p53* plays a critical role in regulating palatogenesis. (A-C) RT-qPCR analysis of *p53, Runx2* and *Sp7* expression in cells isolated from palatal region of control mice 3 days after transfection with *p53* siRNA. Asterisk indicates P < 0.05. (D-E) Histological analysis of samples treated with 10% DMSO in corn oil at E10.5, E12.5 and E14.5. Asterisk in E indicates cleft palate observed in *Wnt1-Cre;Kdm6b^fl/fl^* mouse. Scale bar: 200 µm. (F) p53 protein in the palatal region quantified using western blot. Figure supplement 5-source data 1 for figure supplement 5A Figure supplement 5-source data 2 for figure supplement 5B Figure supplement 5-source data 3 for figure supplement 5C Figure supplement 5-source data 4 for figure supplement 5F Figure supplement 5-source data 5 for figure supplement 5F Figure supplement 5-source data 6 for figure supplement 5F Figure supplement 5-source data 7 for figure supplement 5F Figure supplement 5-source data 8 for figure supplement 5F

**Figure 7-figure supplement 6.**
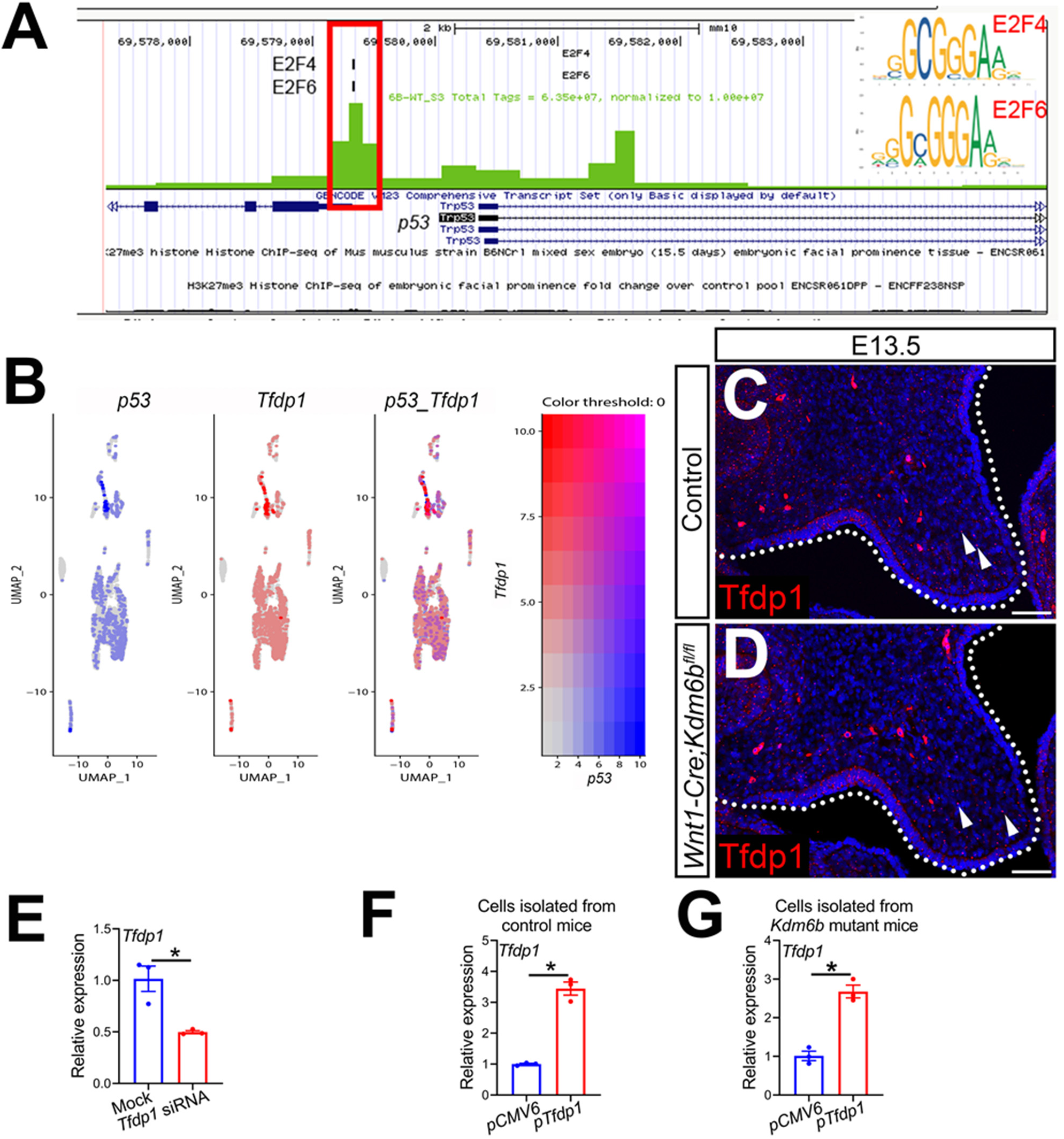
Kdm6b and transcription factors are involved in regulating *p53*. (A) ATAC-seq analysis suggests that the promoter of *p53* is accessible to transcription factors E2f4 and E2f6. (B) Co-expression of *p53* and *Tfdp1*in the palate region at E13.5 using published scRNA-seq analysis (GEO: GSE155928). (C-D) Immunostaining of Tfdp1 in the palatal region of control and *Kdm6b* mutant mice. Arrows indicate representative Tfdp1+ cells. Scale bar: 50 µm. (E) RT-qPCR analysis of *Tfdp1* expression in cells isolated from palatal region of control mice 3 days after transfection with *Tfdp1* siRNA. Asterisk indicates P < 0.05. (F-G) RT-qPCR analysis of *Tfdp1* expression in palatal mesenchymal cells isolated from control and *Kdm6b* mutant mice after transfection with *Tfdp1* overexpressing plasmid. F represents the result using cells isolated from control mice and G represents the result using cells isolated from *Wnt1-Cre;Kdm6b^fl/fl^* mice. Asterisks indicate P < 0.05. Figure supplement 6-source data 1 for figure supplement 6E Figure supplement 6-source data 2 for figure supplement 6F Figure supplement 6-source data 3 for figure supplement 6G

## Supplementary Tables

**Table S1.**

| Antibodies | Vendor | Cat No. | Dilution |
| --- | --- | --- | --- |
| myosin heavy chain (MHC) | DSHB | P13538 | 1:10 |
| Histone H3 tri methyl K27 (H3K27me3) | Cell signaling | 9733s | 1:200 |
| Phospho-Histone H2A.X (Ser139) | Cell signaling | 9718s | 1:200 |
| Dp1 | Abcam | ab124678 | 1:100 |
| Ezh2 | Cell Signaling | 5246s | 1:200 |
| Alexa Fluor 568 | Invitrogen | A-11011 | 1:200 |
| Alexa Fluor 488 | Invitrogen | A-32931 | 1:200 |

**Table S2.**
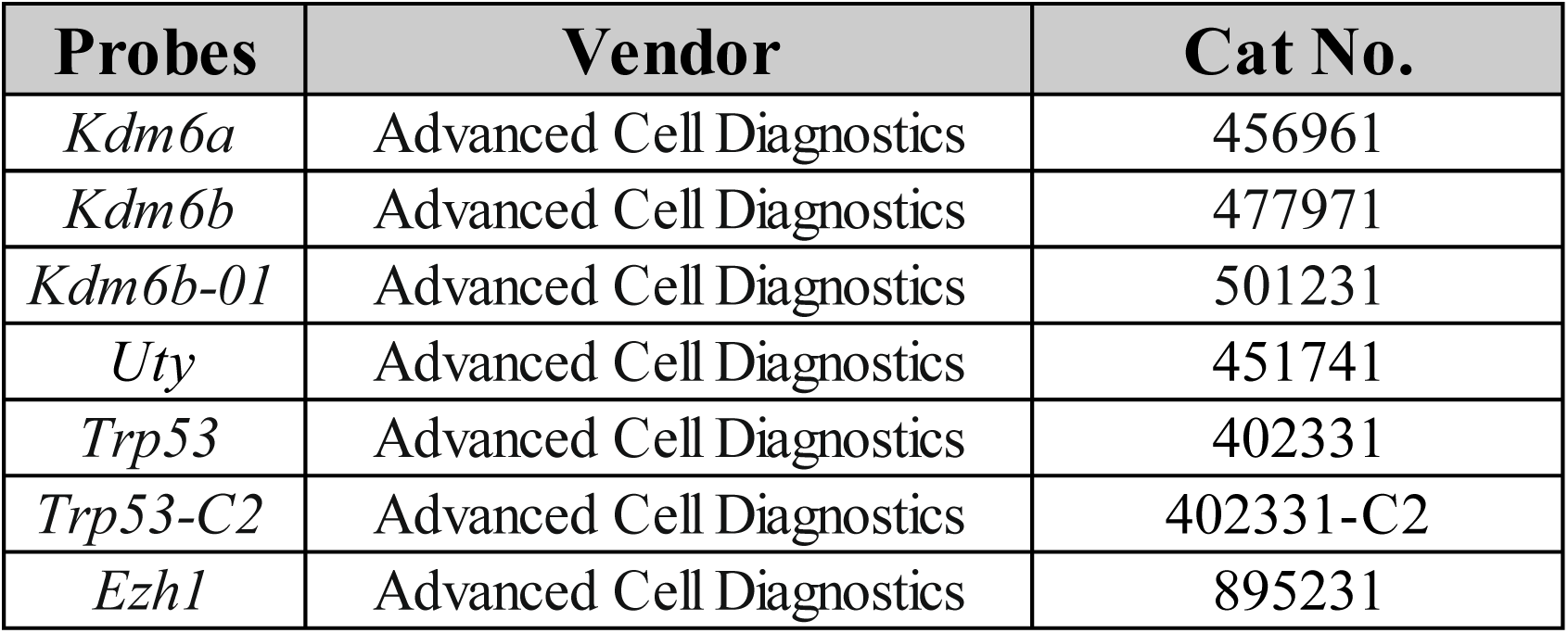

**Table S3.**
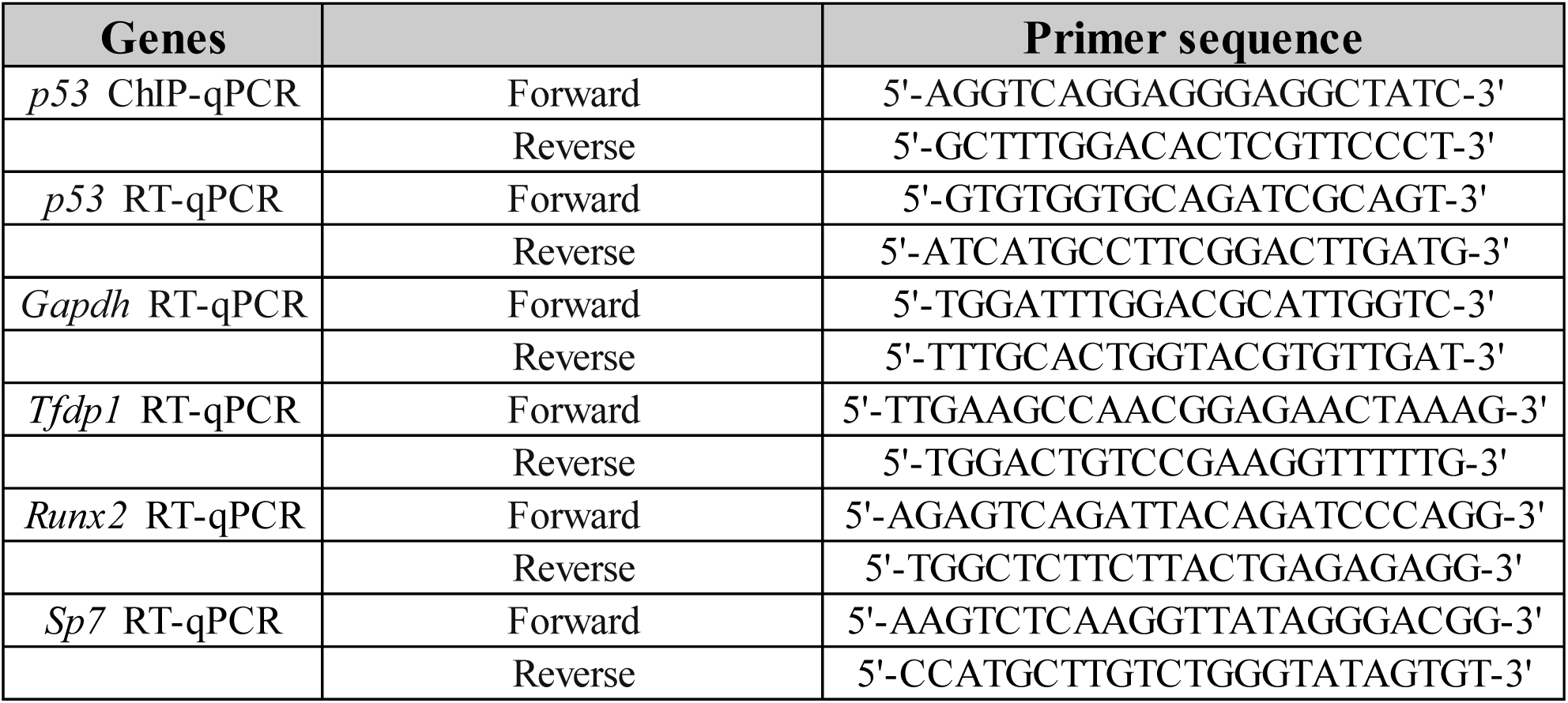

**Table S4.**

| Antibodies | Vendor | Cat No. | Dilution |
| --- | --- | --- | --- |
| Histone H3 tri methyl K27 (H3K27me3) | Cell signaling | 9733s | 1:1000 |
| Dp1 | Abcam | ab124678 | 1:1000 |
| Ezh1 | Abcam | ab189833 | 1:1000 |
| Ezh2 | Cell Signaling | 5246s | 1:2000 |
| Kdm6a | Abcam | ab36938 | 1:1000 |
| Kdm6b (C-term) | Abcepta | AP1022b | 1:1000 |
| Kdm6b (N-term) | Abcepta | AP1022a | 1:1000 |
| p53 | Santa Cruz | sc-126 | 1:1000 |
| Histone H3 | Cell signaling | 4499s | 1:1000 |
| Beta Actin | Abcam | ab20272 | 1:2000 |
| Rabbit IgG HRP-conjugated antibody | R&D System | HAF008 | 1:2000 |
| Mouse IgG HRP-conjugated antibody | R&D System | HAF007 | 1:2000 |
| HRP, Mouse Anti-Rabbit IgG LCS | IPKine™ | A25022 | 1:2000 |

**Table S5.**
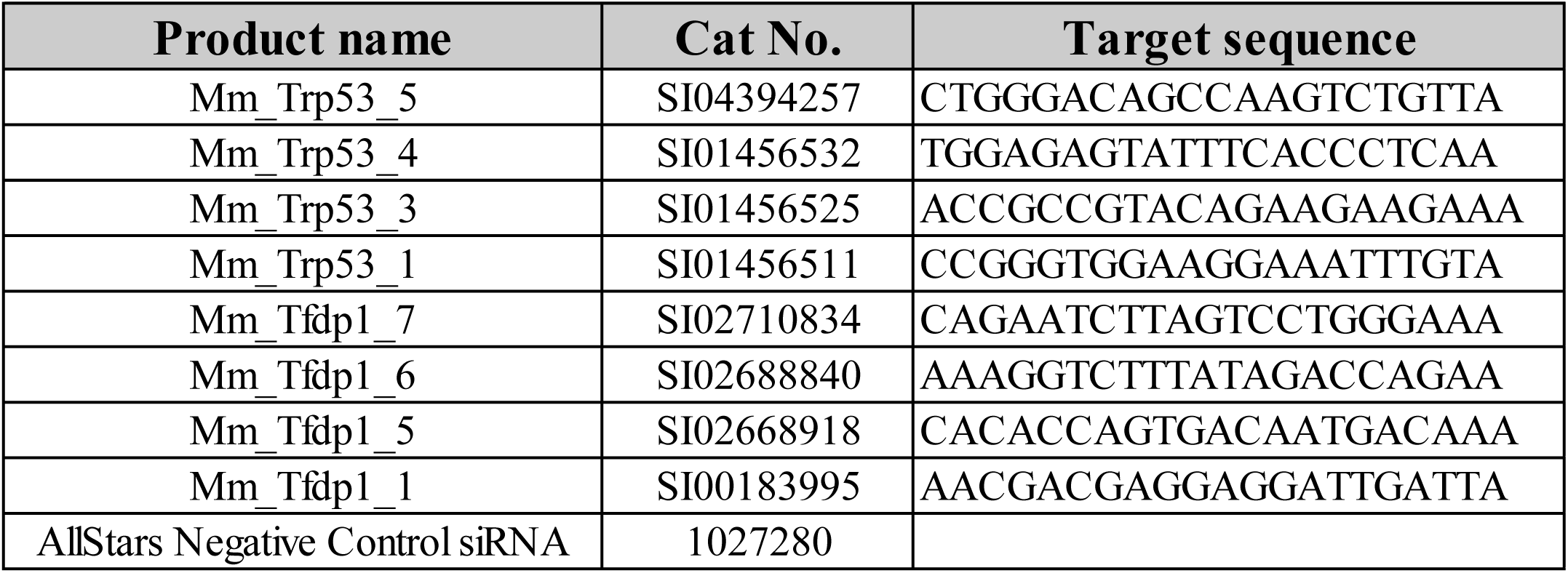

**Table S6.**
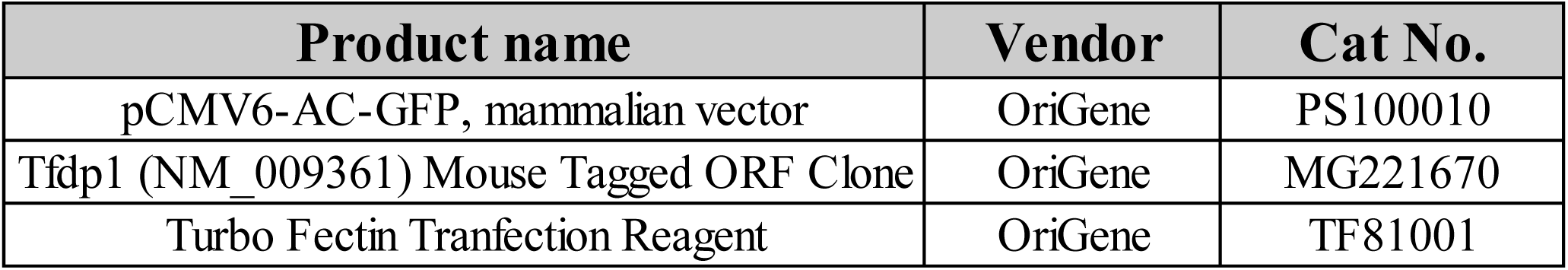

| Product name | Vendor | Cat No. |
| --- | --- | --- |
| pCMV6-AC-GFP, mammalian vector | OriGene | PS100010 |
| Tfdp1 (NM_009361) Mouse Tagged ORF Clone | OriGene | MG221670 |
| Turbo Fectin Transfection Reagent | OriGene | TF81001 |

**Table S7.**

| Pre-alignment qa/qc |  |  |  |  |  |
| --- | --- | --- | --- | --- | --- |
| Sample name | Total reads | Read length | Avg. read quality | % N | % GC |
| 6B-WT_S3 | 117089731 | 76 | 33.4671 | 0.046936 | 44.9143 |

Table S7
| Post-alignment qa/qc |  |  |  |  |  |  |  |  |  |  |  |  |  |  |  |  |  |  |
| --- | --- | --- | --- | --- | --- | --- | --- | --- | --- | --- | --- | --- | --- | --- | --- | --- | --- | --- |
| Sample name | Total reads | Total alignments | Aligned | Total unaligned | Unaligned | Total unique singleton | Unique singleton | Total unique paired | Unique paired | Total non-unique paired | Non-unique paired | Total non-unique singleton | Non-unique singleton | Coverage | Avg. coverage depth | Avg. length | Avg. quality | %GC |
| 6B-WT_S3 | 117089731 | 226531231 | 97.26489422 | 3202528 | 2.735106 | 1951709 | 1.666849 | 1.11E+08 | 94.96082 | 701598 | 0.599197 | 44522 | 0.038024 | 80.3266 | 7.09937 | 75.8895 | 33.6187 | 45.1441 |

## Source Data Files

Figure 2-Source data 1 for figure 2C

Figure 2-Source data 2 for figure 2H

Figure 2-source data 3 for figure 2U

Figure 3-Source data 1 for figure 3H

Figure 3-Source data 2 for figure 3M

Figure 3-Source data 3 for figure 3R

Figure 4-Source data 1 for figure 4C

Figure 4-Source data 2 for figure 4F

Figure 5-source data 1 for figure 5E

Figure 5-source data 2 for figure 5E

Figure 5-source data 3 for figure 5E

Figure 5-source data 4 for figure 5E

Figure 5-source data 5 for figure 5J

Figure 5-source data 6 for figure 5J

Figure 5-source data 7 for figure 5J

Figure 5-source data 8 for figure 5O

Figure 5-source data 9 for figure 5O

Figure 5-source data 10 for figure 5O

Figure 5-source data 11 for figure 5V

Figure 5-source data 12 for figure 5V

Figure 5-source data 13 for figure 5V

Figure 5-source data 14 for figure 5V

Figure 7-source data 1 for figure 7A

Figure 7-source data 2 for figure 7H

Figure 7-source data 3 for figure 7J

Figure 7-source data 4 for figure 7N

Figure 7-source data 5 for figure 7O

Figure 7-source data 6 for figure 7P

Figure 7-source data 7 for figure 7Q

Figure 7-source data 8 for figure 7Q

Figure 7-source data 9 for figure 7Q

Figure 7-source data 10 for figure 7Q

Figure 7-source data 11 for figure 7Q

Figure supplement 1-source data 1 for figure supplement 1M

Figure supplement 1-source data 2 for figure supplement 1M

Figure supplement 1-source data 3 for figure supplement 1M

Figure supplement 1-source data 4 for figure supplement 1N

Figure supplement 1-source data 5 for figure supplement 1N

Figure supplement 1-source data 6 for figure supplement 1N

Figure supplement 5-source data 1 for figure supplement 5A

Figure supplement 5-source data 2 for figure supplement 5B

Figure supplement 5-source data 3 for figure supplement 5C

Figure supplement 5-source data 4 for figure supplement 5F

Figure supplement 5-source data 5 for figure supplement 5F

Figure supplement 5-source data 6 for figure supplement 5F

Figure supplement 5-source data 7 for figure supplement 5F

Figure supplement 5-source data 8 for figure supplement 5F

Figure supplement 6-source data 1 for figure supplement 6E

Figure supplement 6-source data 2 for figure supplement 6F

Figure supplement 6-source data 3 for figure supplement 6G

## References

Arya AK, El-Fert A, Devling T, Eccles RM, Aslam MA, Rubbi CP, Vlatkovic N, Fenwick J, Lloyd BH, Sibson DR et al. 2010. Nutlin-3, the small-molecule inhibitor of MDM2, promotes senescence and radiosensitises laryngeal carcinoma cells harbouring wild-type p53. Br J Cancer 103: 186–195. https://doi.org/10.1038/sj.bjc.6605739

Bannister AJ, Kouzarides T. 2011. Regulation of chromatin by histone modifications. Cell Res 21: 381–395. https://doi.org/10.1038/cr.2011.22

Bernadotte A, Mikhelson VM, Spivak IM. 2016. Markers of cellular senescence. Telomere shortening as a marker of cellular senescence. Aging-Us 8: 3–11. https://doi.org/10.18632/aging.100871

Blagoev KB. 2009. Cell proliferation in the presence of telomerase. PLoS One 4: e4622. https://doi.org/10.1371/journal.pone.0004622

Bowen ME, Attardi LD. 2019. The role of p53 in developmental syndromes. J Mol Cell Biol 11: 200–211. https://doi.org/10.1093/jmcb/mjy087

Bowen ME, McClendon J, Long HK, Sorayya A, Van Nostrand JL, Wysocka J, Attardi LD. 2019. The Spatiotemporal Pattern and Intensity of p53 Activation Dictates Phenotypic Diversity in p53-Driven Developmental Syndromes. Dev Cell 50: 212–228 e216. https://doi.org/10.1016/j.devcel.2019.05.015

Bruneau BG, Koseki H, Strome S, Torres-Padilla ME. 2019. Chromatin and epigenetics in development: a Special Issue. Development 146. https://doi.org/10.1242/dev.185025

Buenrostro JD, Wu B, Chang HY, Greenleaf WJ. 2015. ATAC-seq: A Method for Assaying Chromatin Accessibility Genome-Wide. Curr Protoc Mol Biol 109: 21 29 21-21 29 29. https://doi.org/10.1002/0471142727.mb2129s109

Bush JO, Jiang R. 2012. Palatogenesis: morphogenetic and molecular mechanisms of secondary palate development. Development 139: 231–243. https://doi.org/10.1242/dev.067082

Cardiff RD, Miller CH, Munn RJ. 2014. Manual hematoxylin and eosin staining of mouse tissue sections. Cold Spring Harb Protoc 2014: 655–658. https://doi.org/10.1101/pdb.prot073411

Chai Y, Maxson RE, Jr. 2006. Recent advances in craniofacial morphogenesis. Dev Dyn 235: 2353–2375. https://doi.org/10.1002/dvdy.20833

Chandler RL, Magnuson T. 2016. The SWI/SNF BAF-A complex is essential for neural crest development. Dev Biol 411: 15–24. https://doi.org/10.1016/j.ydbio.2016.01.015

Chandrasekharan D, Ramanathan A. 2014. Identification of a novel heterozygous truncation mutation in exon 1 of ARHGAP29 in an Indian subject with nonsyndromic cleft lip with cleft palate. Eur J Dent 8: 528–532. https://doi.org/10.4103/1305-7456.143637

Cobourne MT, Xavier GM, Depew M, Hagan L, Sealby J, Webster Z, Sharpe PT. 2009. Sonic hedgehog signalling inhibits palatogenesis and arrests tooth development in a mouse model of the nevoid basal cell carcinoma syndrome. Dev Biol 331: 38–49. https://doi.org/10.1016/j.ydbio.2009.04.021

Cordero DR, Brugmann S, Chu Y, Bajpai R, Jame M, Helms JA. 2011. Cranial neural crest cells on the move: their roles in craniofacial development. Am J Med Genet A 155A: 270–279. https://doi.org/10.1002/ajmg.a.33702

den Broeder MJ, Ballangby J, Kamminga LM, Alestrom P, Legler J, Lindeman LC, Kamstra JH. 2020. Inhibition of methyltransferase activity of enhancer of zeste 2 leads to enhanced lipid accumulation and altered chromatin status in zebrafish. Epigenetics Chromatin 13: 5. https://doi.org/10.1186/s13072-020-0329-y

Dimitrova E, Turberfield AH, Klose RJ. 2015. Histone demethylases in chromatin biology and beyond. EMBO Rep 16: 1620–1639. https://doi.org/10.15252/embr.201541113

Dixon MJ, Marazita ML, Beaty TH, Murray JC. 2011. Cleft lip and palate: understanding genetic and environmental influences. Nat Rev Genet 12: 167–178. https://doi.org/10.1038/nrg2933

Flavahan WA, Gaskell E, Bernstein BE. 2017. Epigenetic plasticity and the hallmarks of cancer. Science 357. https://doi.org/10.1126/science.aal2380

Gokbuget D, Blelloch R. 2019. Epigenetic control of transcriptional regulation in pluripotency and early differentiation. Development 146. https://doi.org/10.1242/dev.164772

Grosshans BL, Grotsch H, Mukhopadhyay D, Fernandez IM, Pfannstiel J, Idrissi FZ, Lechner J, Riezman H, Geli MI. 2006. TEDS site phosphorylation of the yeast myosins I is required for ligand-induced but not for constitutive endocytosis of the G protein-coupled receptor Ste2p. Journal of Biological Chemistry 281: 11104–11114. https://doi.org/10.1074/jbc.M508933200

Gurrion C, Uriostegui M, Zurita M. 2017. Heterochromatin Reduction Correlates with the Increase of the KDM4B and KDM6A Demethylases and the Expression of Pericentromeric DNA during the Acquisition of a Transformed Phenotype. J Cancer 8: 2866–2875. https://doi.org/10.7150/jca.19477

Han X, Feng J, Guo T, Loh YE, Yuan Y, Ho TV, Cho CK, Li J, Jing J, Janeckova E et al. 2021. Runx2-Twist1 interaction coordinates cranial neural crest guidance of soft palate myogenesis. Elife 10. https://doi.org/10.7554/eLife.62387

Hanna CW, Demond H, Kelsey G. 2018. Epigenetic regulation in development: is the mouse a good model for the human? Hum Reprod Update 24: 556–576. https://doi.org/10.1093/humupd/dmy021

He F, Chen Y. 2012. Wnt signaling in lip and palate development. Front Oral Biol 16: 81–90. https://doi.org/10.1159/000337619

Heinz S, Benner C, Spann N, Bertolino E, Lin YC, Laslo P, Cheng JX, Murre C, Singh H, Glass CK. 2010. Simple combinations of lineage-determining transcription factors prime cis-regulatory elements required for macrophage and B cell identities. Mol Cell 38: 576–589. https://doi.org/10.1016/j.molcel.2010.05.004

Henckel A, Toth S, Arnaud P. 2007. Early mouse embryo development: could epigenetics influence cell fate determination? Bioessays 29: 520–524. https://doi.org/10.1002/bies.20591

Hobbs CA, Chowdhury S, Cleves MA, Erickson S, MacLeod SL, Shaw GM, Shete S, Witte JS, Tycko B. 2014. Genetic epidemiology and nonsyndromic structural birth defects: from candidate genes to epigenetics. JAMA Pediatr 168: 371–377. https://doi.org/10.1001/jamapediatrics.2013.4858

Hu N, Strobl-Mazzulla P, Sauka-Spengler T, Bronner ME. 2012. DNA methyltransferase3A as a molecular switch mediating the neural tube-to-neural crest fate transition. Genes Dev 26: 2380–2385. https://doi.org/10.1101/gad.198747.112

Hu N, Strobl-Mazzulla PH, Bronner ME. 2014. Epigenetic regulation in neural crest development. Dev Biol 396: 159–168. https://doi.org/10.1016/j.ydbio.2014.09.034

Jambhekar A, Dhall A, Shi Y. 2019. Roles and regulation of histone methylation in animal development. Nat Rev Mol Cell Biol 20: 625–641. https://doi.org/10.1038/s41580-019-0151-1

Jiang W, Wang J, Zhang Y. 2013. Histone H3K27me3 demethylases KDM6A and KDM6B modulate definitive endoderm differentiation from human ESCs by regulating WNT signaling pathway. Cell Res 23: 122–130. https://doi.org/10.1038/cr.2012.119

Jones NC, Lynn ML, Gaudenz K, Sakai D, Aoto K, Rey JP, Glynn EF, Ellington L, Du C, Dixon J et al. 2008. Prevention of the neurocristopathy Treacher Collins syndrome through inhibition of p53 function. Nature Medicine 14: 125–133. https://doi.org/10.1038/nm1725

Kang S, Chovatiya G, Tumbar T. 2019. Epigenetic control in skin development, homeostasis and injury repair. Exp Dermatol 28: 453–463. https://doi.org/10.1111/exd.13872

Kim H, Kim D, Choi SA, Kim CR, Oh SK, Pyo KE, Kim J, Lee SH, Yoon JB, Zhang Y et al. 2018. KDM3A histone demethylase functions as an essential factor for activation of JAK2-STAT3 signaling pathway. Proc Natl Acad Sci U S A 115: 11766–11771. https://doi.org/10.1073/pnas.1805662115

Kim KC, Friso S, Choi SW. 2009. DNA methylation, an epigenetic mechanism connecting folate to healthy embryonic development and aging. J Nutr Biochem 20: 917–926. https://doi.org/10.1016/j.jnutbio.2009.06.008

Kim WT, Kim H, Katanaev VL, Joon Lee S, Ishitani T, Cha B, Han JK, Jho EH. 2012. Dual functions of DP1 promote biphasic Wnt-on and Wnt-off states during anteroposterior neural patterning. EMBO J 31: 3384–3397. https://doi.org/10.1038/emboj.2012.181

Klemm SL, Shipony Z, Greenleaf WJ. 2019. Chromatin accessibility and the regulatory epigenome. Nat Rev Genet 20: 207–220. https://doi.org/10.1038/s41576-018-0089-8

Kohn MJ, Bronson RT, Harlow E, Dyson NJ, Yamasaki L. 2003. Dp1 is required for extra-embryonic development. Development 130: 1295–1305. https://doi.org/10.1242/dev.00355

Lee YH, Saint-Jeannet JP. 2011. Sox9 function in craniofacial development and disease. Genesis 49: 200–208. https://doi.org/10.1002/dvg.20717

Leoyklang P, Suphapeetiporn K, Siriwan P, Desudchit T, Chaowanapanja P, Gahl WA, Shotelersuk V. 2007. Heterozygous nonsense mutation SATB2 associated with cleft palate, osteoporosis, and cognitive defects. Hum Mutat 28: 732–738. https://doi.org/10.1002/humu.20515

Lessard JA, Crabtree GR. 2010. Chromatin regulatory mechanisms in pluripotency. Annu Rev Cell Dev Biol 26: 503–532. https://doi.org/10.1146/annurev-cellbio-051809-102012

Levi G, Mantero S, Barbieri O, Cantatore D, Paleari L, Beverdam A, Genova F, Robert B, Merlo GR. 2006. Msx1 and Dlx5 act independently in development of craniofacial skeleton, but converge on the regulation of Bmp signaling in palate formation. Mech Dev 123: 3–16. https://doi.org/10.1016/j.mod.2005.10.007

Li H. 2013. Aligning sequence reads, clone sequences and assembly contigs with BWA-MEM. *arXiv:13033997v2 [q-bioGN]*.

Li Y, Stockton ME, Bhuiyan I, Eisinger BE, Gao Y, Miller JL, Bhattacharyya A, Zhao XY. 2016. MDM2 inhibition rescues neurogenic and cognitive deficits in a mouse model of fragile X syndrome. Sci Transl Med 8. https://doi.org/10.1126/scitranslmed.aad9370

Lindgren AM, Hoyos T, Talkowski ME, Hanscom C, Blumenthal I, Chiang C, Ernst C, Pereira S, Ordulu Z, Clericuzio C et al. 2013. Haploinsufficiency of KDM6A is associated with severe psychomotor retardation, global growth restriction, seizures and cleft palate. Hum Genet 132: 537–552. https://doi.org/10.1007/s00439-013-1263-x

Lu S, Yang Y, Du Y, Cao LL, Li M, Shen C, Hou T, Zhao Y, Wang H, Deng D et al. 2015. The transcription factor c-Fos coordinates with histone lysine-specific demethylase 2A to activate the expression of cyclooxygenase-2. Oncotarget 6: 34704–34717. https://doi.org/10.18632/oncotarget.5474

Madisen L, Zwingman TA, Sunkin SM, Oh SW, Zariwala HA, Gu H, Ng LL, Palmiter RD, Hawrylycz MJ, Jones AR et al. 2010. A robust and high-throughput Cre reporting and characterization system for the whole mouse brain. Nat Neurosci 13: 133–140. https://doi.org/10.1038/nn.2467

Manna S, Kim JK, Bauge C, Cam M, Zhao Y, Shetty J, Vacchio MS, Castro E, Tran B, Tessarollo L et al. 2015. Histone H3 Lysine 27 demethylases Jmjd3 and Utx are required for T-cell differentiation. Nat Commun 6: 8152. https://doi.org/10.1038/ncomms9152

Marino S, Vooijs M, van Der Gulden H, Jonkers J, Berns A. 2000. Induction of medulloblastomas in p53-null mutant mice by somatic inactivation of Rb in the external granular layer cells of the cerebellum. Genes Dev 14: 994–1004.

Miermont A, Antolovic V, Lenn T, Nichols JME, Millward LJ, Chubb JR. 2019. The fate of cells undergoing spontaneous DNA damage during development. Development 146. https://doi.org/10.1242/dev.174268

Mijit M, Caracciolo V, Melillo A, Amicarelli F, Giordano A. 2020. Role of p53 in the Regulation of Cellular Senescence. Biomolecules 10. https://doi.org/10.3390/biom10030420

Molina-Serrano D, Kyriakou D, Kirmizis A. 2019. Histone Modifications as an Intersection Between Diet and Longevity. Front Genet 10: 192. https://doi.org/10.3389/fgene.2019.00192

Moller M, Schotanus K, Soyer JL, Haueisen J, Happ K, Stralucke M, Happel P, Smith KM, Connolly LR, Freitag M et al. 2019. Destabilization of chromosome structure by histone H3 lysine 27 methylation. PLoS Genet 15: e1008093. https://doi.org/10.1371/journal.pgen.1008093

Nakamura Y, Yamamoto K, He XJ, Otsuki B, Kim Y, Murao H, Soeda T, Tsumaki N, Deng JM, Zhang ZP et al. 2011. Wwp2 is essential for palatogenesis mediated by the interaction between Sox9 and mediator subunit 25. Nature Communications 2. https://doi.org/10.1038/ncomms1242

Nelms BL, Labosky PA. 2010. in Transcriptional Control of Neural Crest Development, San Rafael (CA).

Noden DM. 1983. The role of the neural crest in patterning of avian cranial skeletal, connective, and muscle tissues. Dev Biol 96: 144–165. https://doi.org/10.1016/0012-1606(83)90318-4

Noden DM. 1991. Cell movements and control of patterned tissue assembly during craniofacial development. J Craniofac Genet Dev Biol 11: 192–213.

Parada C, Chai Y. 2012. Roles of BMP signaling pathway in lip and palate development. Front Oral Biol 16: 60–70. https://doi.org/10.1159/000337617

Pediconi N, Salerno D, Lupacchini L, Angrisani A, Peruzzi G, De Smaele E, Levrero M, Belloni L. 2019. EZH2, JMJD3, and UTX epigenetically regulate hepatic plasticity inducing retro-differentiation and proliferation of liver cells. Cell Death Dis 10: 518. https://doi.org/10.1038/s41419-019-1755-2

Reynolds K, Kumari P, Sepulveda Rincon L, Gu R, Ji Y, Kumar S, Zhou CJ. 2019. Wnt signaling in orofacial clefts: crosstalk, pathogenesis and models. Dis Model Mech 12. https://doi.org/10.1242/dmm.037051

Rigueur D, Lyons KM. 2014. Whole-mount skeletal staining. Methods Mol Biol 1130: 113–121. https://doi.org/10.1007/978-1-62703-989-5_9

Rinon A, Molchadsky A, Nathan E, Yovel G, Rotter V, Sarig R, Tzahor E. 2011. p53 coordinates cranial neural crest cell growth and epithelial-mesenchymal transition/delamination processes. Development 138: 1827–1838. https://doi.org/10.1242/dev.053645

Roessler E, Velez JI, Zhou N, Muenke M. 2012. Utilizing prospective sequence analysis of SHH, ZIC2, SIX3 and TGIF in holoprosencephaly probands to describe the parameters limiting the observed frequency of mutant gene x gene interactions. Mol Genet Metab 105: 658–664. https://doi.org/10.1016/j.ymgme.2012.01.005

Ruijtenberg S, van den Heuvel S. 2016. Coordinating cell proliferation and differentiation: Antagonism between cell cycle regulators and cell type-specific gene expression. Cell Cycle 15: 196–212. https://doi.org/10.1080/15384101.2015.1120925

Satokata I, Maas R. 1994. Msx1 deficient mice exhibit cleft palate and abnormalities of craniofacial and tooth development. Nat Genet 6: 348–356. https://doi.org/10.1038/ng0494-348

Schwarz D, Varum S, Zemke M, Scholer A, Baggiolini A, Draganova K, Koseki H, Schubeler D, Sommer L. 2014. Ezh2 is required for neural crest-derived cartilage and bone formation. Development 141: 867–877. https://doi.org/10.1242/dev.094342

Seelan RS, Mukhopadhyay P, Pisano MM, Greene RM. 2012. Developmental epigenetics of the murine secondary palate. ILAR J 53: 240–252. https://doi.org/10.1093/ilar.53.3-4.240

Sen R, Lencer E, Geiger EA, Jones KL, Shaikh TH, Artinger KB. 2020. The role of KMT2D and KDM6A in cardiac development: A cross-species analysis in humans, mice, and zebrafish. bioRxiv: 2020.2004.2003.024646. https://doi.org/10.1101/2020.04.03.024646

Shahbazi MN, Jedrusik A, Vuoristo S, Recher G, Hupalowska A, Bolton V, Fogarty NNM, Campbell A, Devito L, Ilic D et al. 2016. Self-organization of the human embryo in the absence of maternal tissues. Nat Cell Biol 18: 700–708. https://doi.org/10.1038/ncb3347

Shen X, Liu Y, Hsu YJ, Fujiwara Y, Kim J, Mao X, Yuan GC, Orkin SH. 2008. EZH1 mediates methylation on histone H3 lysine 27 and complements EZH2 in maintaining stem cell identity and executing pluripotency. Mol Cell 32: 491–502. https://doi.org/10.1016/j.molcel.2008.10.016

Shpargel KB, Starmer J, Wang C, Ge K, Magnuson T. 2017. UTX-guided neural crest function underlies craniofacial features of Kabuki syndrome. Proc Natl Acad Sci U S A 114: E9046–E9055. https://doi.org/10.1073/pnas.1705011114

Smith ZD, Meissner A. 2013. DNA methylation: roles in mammalian development. Nat Rev Genet 14: 204–220. https://doi.org/10.1038/nrg3354

Soares LM, He PC, Chun Y, Suh H, Kim T, Buratowski S. 2017. Determinants of Histone H3K4 Methylation Patterns. Mol Cell 68: 773–785 e776. https://doi.org/10.1016/j.molcel.2017.10.013

Soldatov R, Kaucka M, Kastriti ME, Petersen J, Chontorotzea T, Englmaier L, Akkuratova N, Yang YS, Haring M, Dyachuk V et al. 2019. Spatiotemporal structure of cell fate decisions in murine neural crest. Science 364: 971-+. https://doi.org/10.1126/science.aas9536

Sugii H, Grimaldi A, Li J, Parada C, Vu-Ho T, Feng J, Jing J, Yuan Y, Guo Y, Maeda H et al. 2017. The Dlx5-FGF10 signaling cascade controls cranial neural crest and myoblast interaction during oropharyngeal patterning and development. Development 144: 4037–4045. https://doi.org/10.1242/dev.155176

Tateossian H, Morse S, Simon MM, Dean CH, Brown SD. 2015. Interactions between the otitis media gene, Fbxo11, and p53 in the mouse embryonic lung. Dis Model Mech 8: 1531–1542. https://doi.org/10.1242/dmm.022426

Trainor P, Krumlauf R. 2000. Plasticity in mouse neural crest cells reveals a new patterning role for cranial mesoderm. Nat Cell Biol 2: 96–102. https://doi.org/10.1038/35000051

Wiles ET, Selker EU. 2017. H3K27 methylation: a promiscuous repressive chromatin mark. Curr Opin Genet Dev 43: 31–37. https://doi.org/10.1016/j.gde.2016.11.001

Williams AB, Schumacher B. 2016. p53 in the DNA-Damage-Repair Process. Cold Spring Harb Perspect Med 6. https://doi.org/10.1101/cshperspect.a026070

Wilson S, Filipp FV. 2018. A network of epigenomic and transcriptional cooperation encompassing an epigenomic master regulator in cancer. NPJ Syst Biol Appl 4: 24. https://doi.org/10.1038/s41540-018-0061-4

Wysocka J, Swigut T, Xiao H, Milne TA, Kwon SY, Landry J, Kauer M, Tackett AJ, Chait BT, Badenhorst P et al. 2006. A PHD finger of NURF couples histone H3 lysine 4 trimethylation with chromatin remodelling. Nature 442: 86–90. https://doi.org/10.1038/nature04815

Xu J, Liu H, Lan Y, Aronow BJ, Kalinichenko VV, Jiang R. 2016. A Shh-Foxf-Fgf18-Shh Molecular Circuit Regulating Palate Development. PLoS Genet 12: e1005769. https://doi.org/10.1371/journal.pgen.1005769

Young JI, Slifer S, Hecht JT, Blanton SH. 2021. DNA Methylation Variation Is Identified in Monozygotic Twins Discordant for Non-syndromic Cleft Lip and Palate. Front Cell Dev Biol 9. https://doi.org/10.3389/fcell.2021.656865

Yu L, Gu S, Alappat S, Song Y, Yan M, Zhang X, Zhang G, Jiang Y, Zhang Z, Zhang Y et al. 2005. Shox2-deficient mice exhibit a rare type of incomplete clefting of the secondary palate. Development 132: 4397–4406. https://doi.org/10.1242/dev.02013

Zhang Y, Liu T, Meyer CA, Eeckhoute J, Johnson DS, Bernstein BE, Nusbaum C, Myers RM, Brown M, Li W et al. 2008. Model-based analysis of ChIP-Seq (MACS). Genome Biol 9: R137. https://doi.org/10.1186/gb-2008-9-9-r137

Zhao H, Oka K, Bringas P, Kaartinen V, Chai Y. 2008. TGF-beta type I receptor Alk5 regulates tooth initiation and mandible patterning in a type II receptor-independent manner. Dev Biol 320: 19–29. https://doi.org/10.1016/j.ydbio.2008.03.045

Zoghbi HY, Beaudet AL. 2016. Epigenetics and Human Disease. Cold Spring Harb Perspect Biol 8: a019497. https://doi.org/10.1101/cshperspect.a019497

